# Evaluating horseradish peroxidase-mimic DNAzyme transducer for glucometer readout: roles of various components and their optimization

**DOI:** 10.1101/2025.06.27.661989

**Authors:** Saba Parveen, Arunanshu Talukdar, Mrittika Sengupta, Souradyuti Ghosh

**Affiliations:** Centre for Life Sciences, Mahindra University, Hyderabad, Telangana, India; Interdisciplinary Centre for Nanosensors and Nanomedicine, Mahindra University, Hyderabad, Telangana, India; Department of Geriatric Medicine, Medical College and Hospital Kolkata, Kolkata

**Keywords:** Peroxidase mimic-DNAzyme, Glucometer, Mobile Health, Device repurposing

## Abstract

The Development of decentralized disease detection elements is integral towards ensuring democratized access to healthcare. Personal glucometers, thanks to their low cost, ease of usage, and universal availability, are increasingly being identified as a central tenet of democratized disease detection. Towards this objective, it will first be critical to establish whether glucometers can be integrated with established molecular diagnosis workflows such as enzyme-linked absorbent assay (ELISA) or its analogous techniques such as enzyme-linked aptasorbent assays (ELASA). In ELISA or its analogous assays, horseradish peroxidase (HRP) or its mimics, such as hemin DNAzyme conjugated to biorecognition elements (e.g., antibody or aptamers), serve as the signal transducers. These transducers generate colorimetric or electrochemical signals measurable in centralized instruments such as ELISA readers or potentiostats. To become compatible with ELISA readouts in decentralized settings, the glucometers should therefore be able to quantify the activity of HRP mimics such as hemin DNAzyme. The current work thus explores the possibility of the use of glucometers as a general readout instrument for hemin DNAzyme transducer activity involving potassium ferrocyanide as a novel redox substrate. Using both absorbance and glucometer readout, we systematically investigated the role of buffer, pH, redox mediator, and H_2_O_2_ in achieving optimal readout conditions. It also probed the prospective DNAzyme operational range under these assay conditions. In addition, transferability of optimized assay parameters in other commercial glucometer brands, as well as preliminary suitability in HRP enzyme, was explored. Altogether, our study explores the possibility of using DNAzymes as transducers for glucometers as a democratized alternative to ELISA readers, multimode readers, or potentiostats towards equitable access to disease diagnosis.

## Introduction

Enzyme-linked immunosorbent assay (ELISA) or its analogues, such as enzyme-linked aptasorbent assay (ELASA) are integral to molecular disease diagnosis. First reported by Engvall and Perlmann in 1971^1^, they use HRP or HRP mimics (such as hemin G-quadruplex DNAzyme^2^, CeO_2_ nanoparticle^3^) as the transducer. These enzymes, analogues, or their mimics catalyze the reduction of H_2_O_2_ to H_2_O while simultaneously oxidizing a substrate for a favorable chromogenic, fluorogenic, or electrogenic readout. Alongside ELISA or ELASA, this HRP or its analogue-enabled reporter chemistry has undergone extensive biochemical engineering and demonstrated for nucleic acid detection^4,5^. Similarly, multimeric HRP molecules conjugated to nanoparticles^6^, multiple hemin DNAzymes embedded within DNA nanostructures^7^, or multimodal nanoparticle moieties have been developed for these applications. Among substrates, 2,2’-azino-bis(3-ethylbenzothiazoline-6-sulfonic acid) (ABTS) or 3,3’,5,5’-tetramethylbenzidine (TMB) are used as the primary chromogenic substrates^8,9^. Upon oxidation by HRP or its analogues, such as hemin DNAzyme, ABTS and TMB convert to radical cations, absorbing at specific wavelengths (415 and 450 nm, respectively) and displaying color (green in citrate-phosphate buffer). Additionally, common substrates such as Amplex red or 2′,7′-dichlorodihydrofluorescein (converts to fluorescent resorufin, excitation/emission 570/5855 nm)^10^, luminol (for luminescence, substrate for HRP or hemin DNAzymes^11,12^), TMB (for electrochemical readout for HRP or DNAzyme mimics^5,13^) or recently developed molecules such as phenylenediamine^14^ or formation of fluorescent polyethyleneimine-phenylenediamine copolymer nanoparticles are known^15^. Besides commercial molecular diagnostic applications, these assays have been engineered to detect a plethora of molecular targets such as heavy metal ions^7^, SARS-CoV-2 nucleic acids^4^, carbohydrates^16^, lipids^17^, pathogens^18^, and cancer cells^19^. Despite established substrates and standard operating procedures in these workflows, these methods rely on ELISA readers, multimode readers, or potentiostats for quantitative, semi-quantitative, or qualitative readouts. However, all these instruments are centralized, non-portable, and electricity-intensive. Harnessing these powerful and sensitive assays, therefore, necessitates sample transport to centralized labs, not only causing delay in diagnosis, but also contributing to unsustainable electricity-intensive instrument usage, cost enhancement, and most importantly, restriction of disease detection at resource-constrained settings.

Perhaps the most discussed use of alternative assays that are amenable to resource-constrained settings remains the utilization of the glucometer. An over-the-counter glucometer is one of the most universally accessible, low-cost (at least 1000 times less costly than an ELISA reader or similar instruments), and user-friendly biomedical instruments. It uses glucometer strips that contain a cocktail of glucose oxidase or an analogous glucose-selective enzyme (e.g., glucose dehydrogenase) along with a mediator such as potassium ferricyanide/ferrocyanide redox couple^20^. Upon exposure to glucose, it triggers a redox chemistry catalyzed by the enzymatic oxidation of glucose and subsequent electron transfer between the redox mediator and glucometer strip electrode. This mechanism has been fortuitously utilized by Xiang et al. who conjugated integrase enzyme (replacing the HRP enzyme) to antibodies or aptamers^21^. In this chemistry, sucrose replaces ATBS or TMB at the final step of an ELISA assay. If an antibody-coupled integrase or a similar recognition-transducer conjugate is present, it will convert sucrose to glucose. Upon exposure to a glucometer strip mounted on a glucometer, the presence of glucose will be recognized, essentially helping to draw a quantitative or semi-quantitative conclusion for the ELISA assay. A similar assay has been developed via conjugation of glucoamylase to aptamers, which utilizes amylose as a substrate to generate glucose^22^. Both the invertase and glucoamylase conjugations have been adopted for nucleic acid reporters as well as for aptameric or genosensor applications. However, both these classes of assays require chemical conjugation of invertase and glucoamylase to antibody or nucleic acid recognition molecules.

Careful consideration thus highlights that all the steps of ELISA-like assays can be undertaken using simple apparatus such as pipettes, irrespective of location. However, the final readout step involving centralized instruments such as an ELISA reader, multimode reader, or a potentiostat is the major obstacle in the universal and democratized adoption of ELISA, especially in resource-constrained settings. Although invertase or glucoamylase-based assays constitute a major progress towards this direction, they still require chemical conjugation to antibodies or nucleic acids. Therefore, an innovation that would permit the utilization of a glucometer without chemical conjugation of reporter molecule with invertase or glucoamylase will truly bring a transformative change in rural and mobile healthcare.

We identified that some of the earliest literature studies involving the enzymatic activities of HRP were carried out using potassium ferrocyanide as its substrate^23^. This led us to hypothesize that potassium ferrocyanide (Fe(II)) may as well be a direct substrate for HRP-mimic hemin DNAzyme reactivity in the presence of H_2_O_2_, which will convert ferrocyanide to ferricyanide (Fe(III)). On the other hand, potassium ferricyanide, being the redox mediator through glucose oxidase reaction, will likely be directly measurable by a glucometer via a glucometer strip^20^. Together, this opened up the possibility that the ELISA/ELASA-coupled activity of HRP-mimic hemin DNAzyme in the presence of H_2_O_2_ and ferrocyanide may be directly measurable through a glucometer without any chemical conjugation or additional intermediary redox mediator. In this study, we therefore explored the potential usability of combinations of HRP-mimic hemin DNAzyme, H_2_O_2_, and ferrocyanide/ferricyanide redox couple for glucometer readout. We investigated the role of buffers, pH, H_2_O_2_, potassium ferrocyanide, and DNAzyme concentration to optimize the assay conditions. In addition to the glucometer readout, spectroscopic measurements leveraging the ferricyanide absorbance were carried out to gain insight into the mechanistic aspect of the assay. We also investigated the possible usability of the assay across alternate commercial glucometer brands as well as for HRP itself.

## Materials and Methods

Please see the Supporting Information file.

## Results

### Design of experiments

The premise of this study rests on first identifying whether HRP-mimic G-quadruplex hemin DNAzyme could recognize potassium ferrocyanide as a substrate in the presence of H_2_O_2_. Secondly, it was critical to identify if this reaction would generate a readout in a glucometer (Figure 1A). In addition, suitable buffers, H_2_O_2_ or ferrocyanide concentration ranges remain unidentified for such a reaction. To investigate these variables, we prepared an immobilized DNAzyme model (utilizing EAD2 hemin DNAzyme^24^) that would resemble ELASA or similar experimental setups where DNAzymes have been used as transducers (Figure 1B)^25–29^. A streptavidin magnetic bead-bound biotinylated oligonucleotide was utilized to capture (by base pairing between sequences a and a*) and immobilize the active hemin DNAzyme. Given that hemin itself has oxidation capability^30^, we posited that this experimental assembly would also remove any excess unbound hemin from interfering in our redox study. Before proceeding with any further studies, the activity of the immobilized hemin DNAzyme was always first confirmed through ABTS colorimetry at 12.5 and 50 nM DNAzyme concentrations in CPB buffers. Next, the immobilized DNAzyme was subjected to reactions to identify optimal buffer(s) and pH, suitable H_2_O_2_ and ferrocyanide concentration range, and finally, DNAzyme operational range. K_4_[Fe(CN)_6_] and K_3_[Fe(CN)_6_] have λ_max_ at 220 and 420 nm, respectively, while only ferricyanide generates a glucometer response. Thanks to ferrocyanide’s selective spectroscopic signature absorbance and glucometer readout (Figure S1), our study involved a combined spectrophotometric and glucometer measurement. In parallel, we confirmed that potassium ferrocyanide was undetectable in either absorbance (405 nm) or glucometer (data not shown).

**Figure 1.**
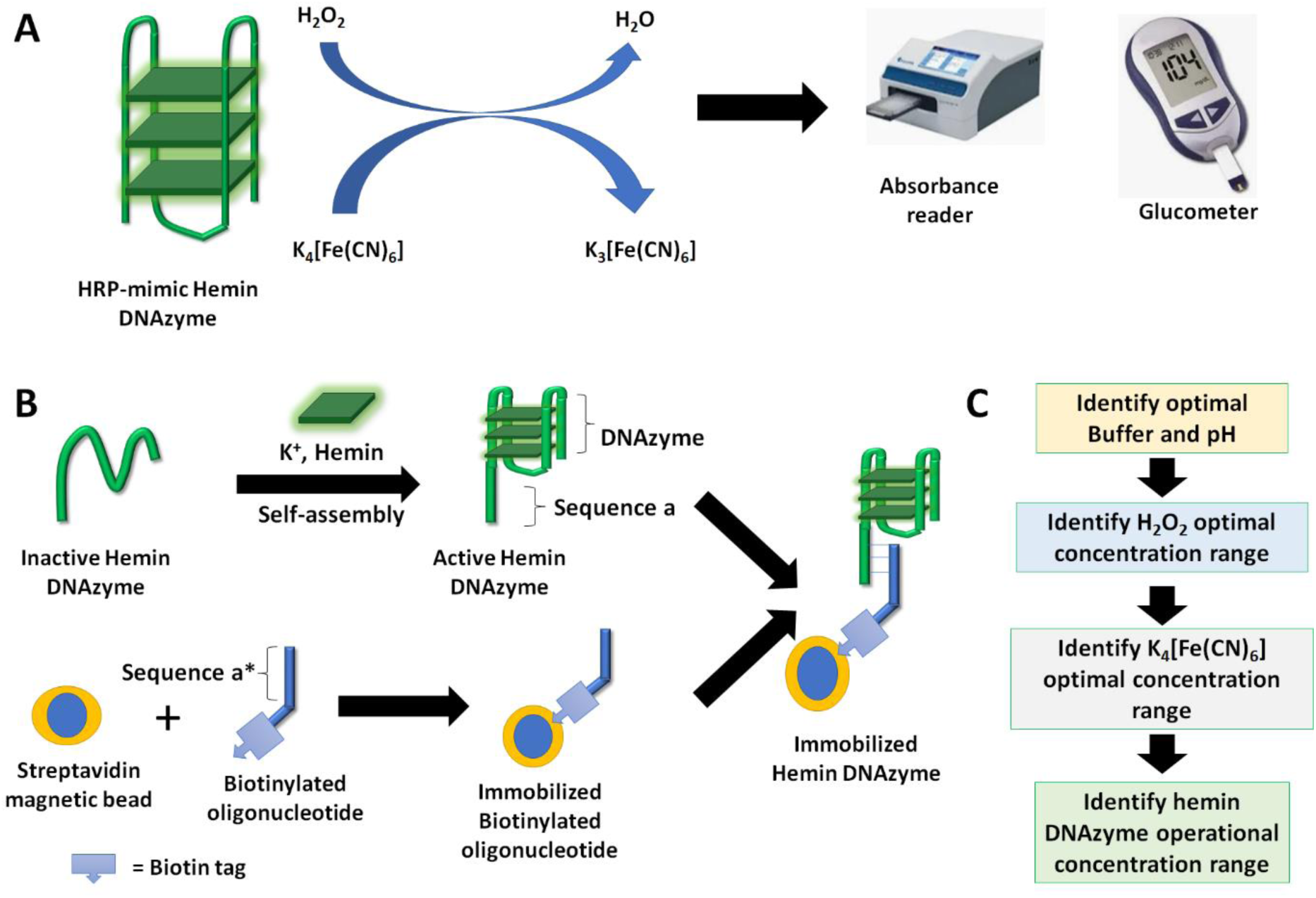
Functional principle and experimental schematics of this study. A, Proposed catalytic oxidation of potassium ferrocyanide by hemin DNAzyme. B, Preparation of magnetic bead immobilized hemin DNAzyme. C, Schematics of workflow for the identification of various operational parameters of this study.

### Hypothesis validation and buffer optimization

First, we wanted to obtain proof-of-concept to validate our hypothesis that DNAzyme would utilize ferrocyanide as a substrate in the presence of H_2_O_2_ (Figure 1C). Next, we wanted to evaluate and identify the buffer conditions(s) and pH where DNAzyme exhibited the highest catalytic efficiency in oxidizing the ferrocyanide substrate. We rationalized that this condition would give a significant difference in absorbance between the presence and absence of DNAzyme. In parallel, we wanted to obtain proof-of-concept to validate our hypothesis that DNAzyme would utilize ferrocyanide as a substrate in the presence of H_2_O_2_. Towards this goal, we investigated different buffers and pH values, such as MES (pH 5.0), sodium acetate (pH 5.0), HEPES (pH 7.2 & pH 8.2), or Tris-HCl (pH 7.6 & pH 8.5), and performed the ferricyanide absorbance measurement (405 nm) as a time course (0-40 min). All these conditions were initially investigated with constant potassium ferrocyanide (10 mM) and H_2_O_2_ (50 mM) concentration for the time being, while the effect of ferrocyanide and H_2_O_2_ was probed later. These experiments were performed with and without DNAzyme (referred to as “DNAzyme+” and “DNAzyme−”, or “DNAzyme present” and “DNAzyme absent”, respectively) and represented in terms of both kinetics and absorbance %fold change between 0-40 min (Figure 2). While the time course helped us understand the DNAzyme behaviour, the difference in 0-40 min absorbance %fold change was indicative of whether the assay can distinguish the DNAzyme presence vs absence. The higher the difference in %fold change between the presence and absence of DNAzyme, the better it would be in differentiating the presence of an analyte compared to its absence.

**Figure 2.**
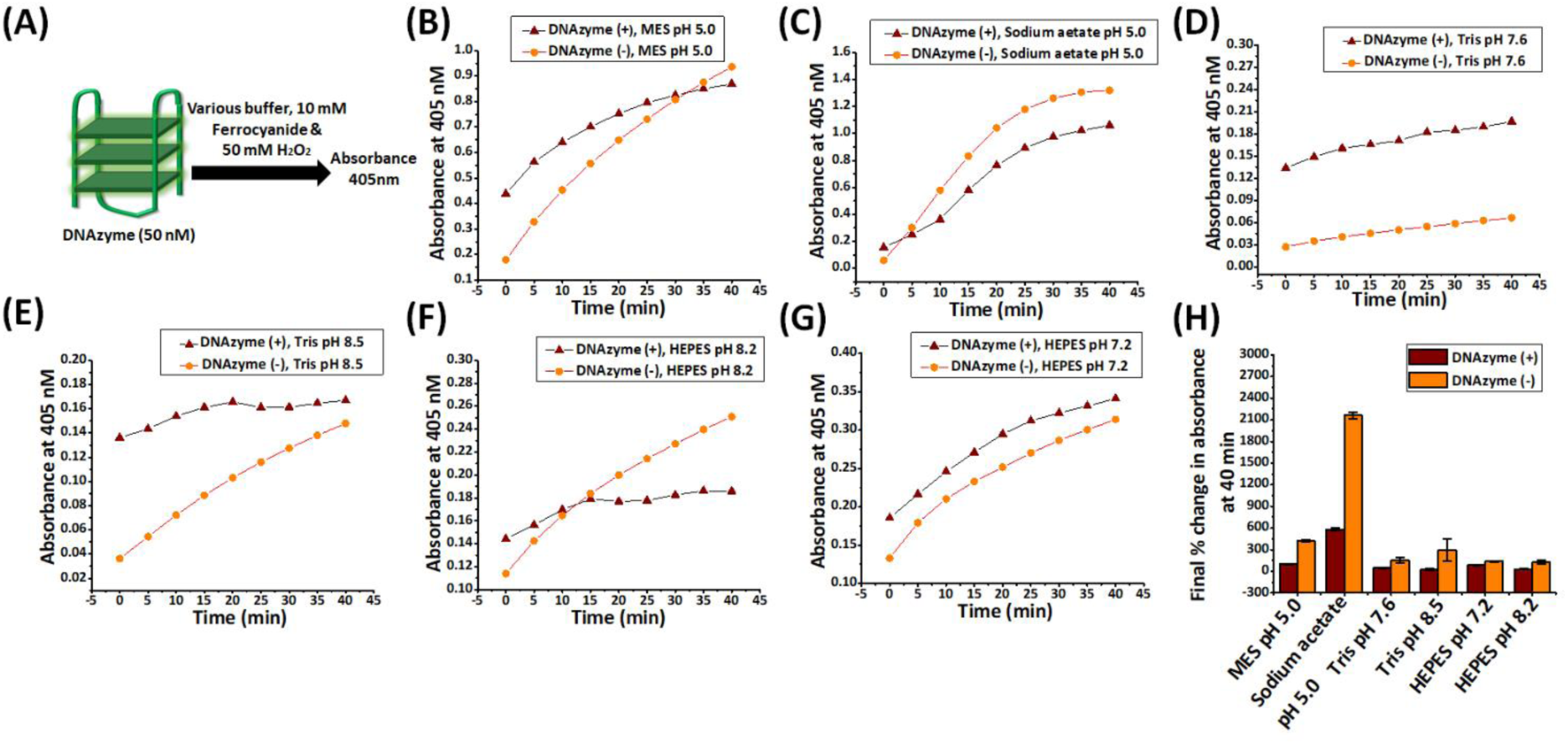
Optimization of buffer and pH for DNAzyme-catalyzed oxidation of potassium ferrocyanide. Panel A represents a schematic of the reaction. Oxidation of potassium ferrocyanide to potassium ferricyanide in the presence or absence of DNAzyme, H_2_O_2_ and buffer MES pH 5 (Panel B), sodium acetate pH 5 (Panel C), Tris pH 7.6 (Panel D), Tris pH 8.2 (Panel E), HEPES pH 7.2 (Panel F), and HEPES pH 8.2 (Panel G). The time-dependent absorbance changes at 405nm indicates the formation of ferricyanide over 40 minutes under different buffers and pH. The reaction mixture contained 10 mM potassium ferrocyanide and 50 mM H_2_O_2_, with and without 50 nM DNAzyme, to assess catalytic activity. Panel H represents the % fold change in the oxidation of ferrocyanide (Fe²⁺) to ferricyanide (Fe³⁺) 0 – 40 minutes in the presence and absence of DNAzyme. Error bars represent standard deviation (n = 3).

All buffer and pH conditions exhibited an absorbance difference in the presence and absence of DNAzyme (Figure 2B-F). A common noticeable trend observed among these conditions was the predominantly linear absorbance increase for the “DNAzyme absent” conditions. In contrast, the absorbance in “DNAzyme present” conditions here, as well as in later studies, were particularly marked by either consistent slower increase throughout the monitoring time (e.g., Figure 2D, 2E) or an early increase followed by plateauing (Figures 2B, 2C, 2F, 2G). This pattern in the difference in absorbance growth curves between DNAzyme presence/absence conditions would remain consistent throughout the rest of the studies reported in this manuscript. The high magnitude of 0 min absorbance readings and 0-10 min readings illustrated that in the presence of DNAzyme, ferrocyanide oxidation occurred rapidly (Figure 2). In some cases, the majority of the oxidation happened even before the first absorbance reading was measured, then reaching a steady state within 15 minutes. This rapid oxidation occurred due to the catalytic activity of DNAzyme, which rapidly sequestered the H_2_O_2_ to oxidize ferrocyanide to ferricyanide. We hypothesized that the plateau at the later parts (15 min onwards) of the assay probably originated from either an irreversible activity loss/reduction of DNAzyme or a (mostly) complete conversion of ferrocyanide to ferricyanide. Since these assays used 10 mM ferrocyanide, its complete consumption would lead to 10 mM ferricyanide, causing an absorbance of approximately 0.8-1.0 (Figure S1B). The “DNAzyme presence” conditions reached these absorbance levels only for MES and sodium acetate, indicating better catalytic activity and complete consumption of ferrocyanide. In contrast, “DNAzyme absent” conditions were marked by the aforementioned characteristic linear absorbance increase, which can be attributed to non-catalytic H_2_O_2_-mediated ferrocyanide oxidation. At the same time, the sodium acetate (pH 5.0) demonstrated the highest fold change differentiating the presence and absence of DNAzyme (Figure 1C and H), presumably due to lower starting absorbance for the latter. In contrast, MES showed the second highest 0-40 min absorbance %fold change between DNAzyme presence/absence conditions (Figure 1B and H). The absorbance behaviour in sodium acetate was consistent when we performed further replicates (data not shown). Between MES and sodium acetate, MES appeared to be a more suitable buffer due to higher initial starting (0 min) absorbance, indicating better catalytic behaviour.

In contrast, “DNAzyme present” reactions conducted at both Tris buffers and HEPES pH 8.2 buffers, the initial (0 min) absorbance magnitude remained lower than their MES or acetate counterparts (albeit higher than “DNAzyme absent” conditions), indicating generally lower enzymatic activity. In fact, for these three buffers (with an exceptional early (0-10 min) burst for HEPES pH 8.2), there were mostly insignificant changes in “DNAzyme present” absorbances throughout the measurement duration (Figure 2D-F). In contrast, after an initial (0 min) greater absorbance for the “DNAzyme present” condition, both DNAzyme present/absent conditions in HEPES pH 7.2 showed linear and almost parallel increases throughout the study (Figure 2H). This could be attributed to an early loss or attenuation of DNAzyme enzymatic activity while simultaneously oxidation of ferrocyanide by H_2_O_2_. The 0-40 min absorbance %fold change between DNAzyme presence/absence conditions for these reactions expectedly reflected this pattern (Figure 2H). Sub-optimal DNAzyme performance in alkaline buffer could be rationalized from existing literature concerning citing ferricyanide oxidizing H_2_O_2_ to oxygen in an alkaline environment (while itself getting reduced to ferrocyanide)^31^. The DNAzyme behaviour is similar to HRP-enabled faster (10^3^-10^5^-fold) ferrocyanide oxidation in an acidic environment compared to the same in HRP absence^23,32^.

Another key repeatable pattern in this experiment and elsewhere in this manuscript was 0-40 min absorbance %fold change for “DNAzyme absent” conditions was usually higher than “DNAzyme present” conditions. As explained above, DNAzyme activity would sequester most of H_2_O_2_ and ferrocyanide within the first few minutes. Non-availability of substrate would then slow down the DNAzyme activity, causing a lesser increase in absorbance during the actual observation time (0-40 min), leading to a diminished absorbance %fold change. Overall, these experiments established our hypothesis about the speculative catalytic role of DNAzyme concerning ferrocyanide as a substrate in its oxidation process. The initial greater absorbance magnitude and consistently higher absorbance %fold change difference in MES pH 5.0 across the replicates (Figure 2B and H) suggest that MES might provide a suitable buffering environment for the assay. While a large difference in DNAzyme presence/absence was observed in sodium acetate and to some extent in HEPES pH 7.2 and Tris pH 7.6, we wanted to further investigate if the results can be replicated through glucometer readouts as well. Due to a lesser fold change and sub-optimal DNAzyme performance, we opted not to continue with Tris pH 8.5 and HEPES pH 8.2.

### Buffer optimization for glucometer reading

In this section, we continued to evaluate and optimize buffer and pH, where the presence and absence of DNAzyme would demonstrate differential performance in terms of glucometer reading (mg/dL). Another goal was to find a suitable time point for capturing the glucometer readout. Towards this objective, we used three different buffers and pH values shortlisted from their performance in absorbance studies (above, Figure 2), namely MES (pH 5.0), HEPES (pH 7.2), and Tris (pH 7.6). The hemin DNAzyme-enabled oxidation of ferrocyanide to ferricyanide was performed under these conditions, taking the glucometer readings from 0 to 40 minutes with and without DNAzyme (Figure 3).

**Figure 3.**
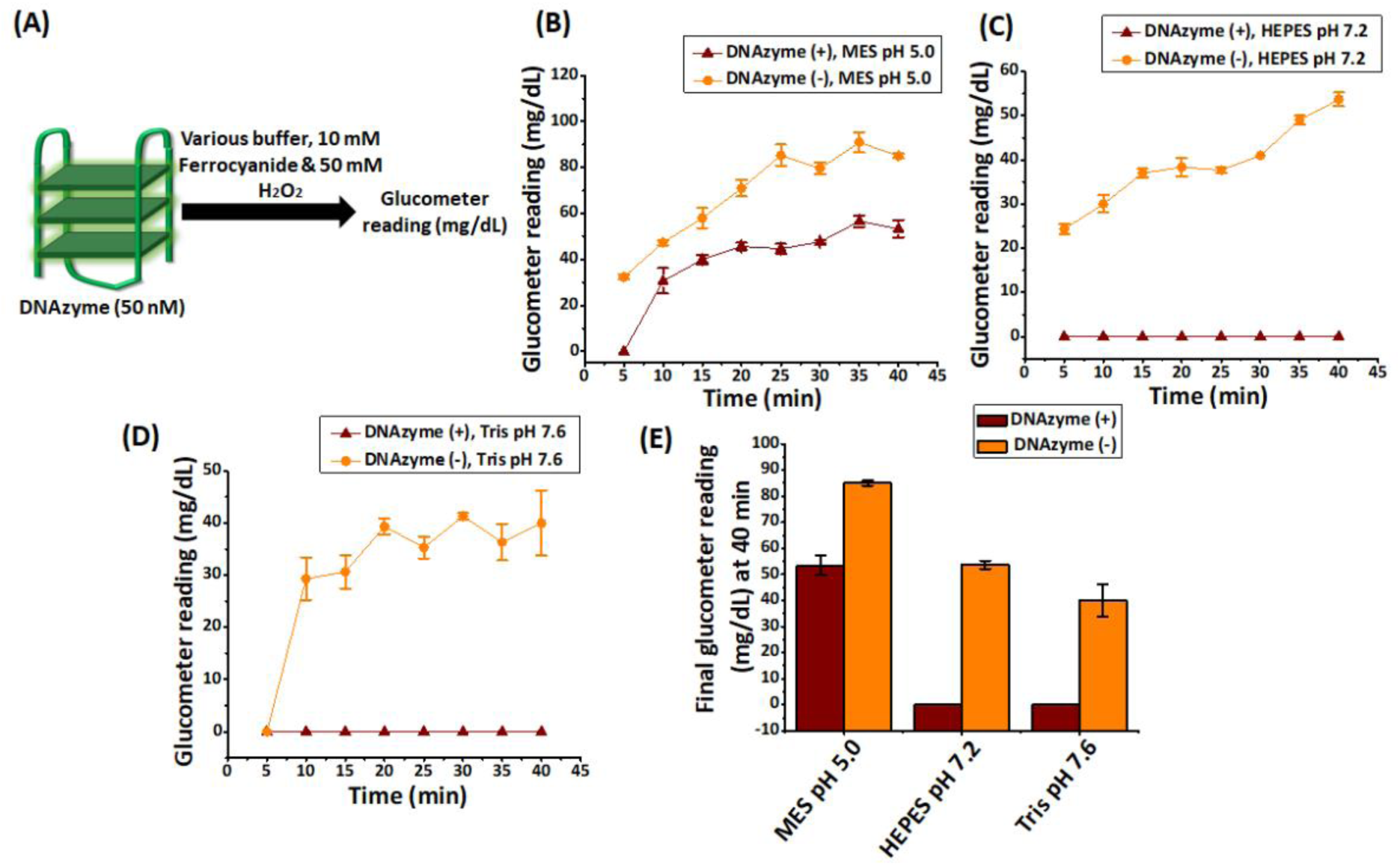
Effect of selected buffer on DNAzyme-catalyzed oxidation of ferrocyanide to ferricyanide monitored by glucometer. Panel A schematic illustration of the experimental setup. Panel B-D represents the time course of glucometer readings (mg/dL) from 0 to 40 minutes, measured at 5-minute intervals, in different buffer systems: MES pH 5.0 (B), HEPES pH 7.2 (C), and Tris pH 7.6 (D) in the presence and absence of DNAzyme. Panel E represent the final glucometer reading at 40 minutes, indicating the extent of ferrocyanide oxidation to ferricyanide in the presence or absence of DNAzyme across the three buffer conditions. Error bars represent standard deviation (n = 3) Experiments were performed with Dr. Morepen GLuco One brand.

In HEPES (pH 7.2) and Tris (pH 7.6) buffers, no measurable glucometer signal in the presence of DNAzyme was observed even after 40 minutes (Figure 3C, 3D, and 3E). In comparison, DNAzyme readings started to appear within 10 min for MES, re-affirming higher catalytic activity in it. In MES, the DNAzyme activity appeared to reach a steady state by 15 min (Figure 3B). In contrast, in the absence of DNAzyme, ferrocyanide oxidation started earlier (within 5 minutes) in MES buffer and continued linearly increasing up to 40 minutes (Figure 3B), indicating a non-enzymatic oxidation process. Additionally, the data implied that the maximum difference in glucometer readout between the presence/absence of DNAzyme would be obtained at 30-40 min. This was consistent with the absorbance data shown in Figure 2B and reproducible across additional biological replicates carried out by us (data not shown).

The complete absence of glucometer readings for “DNAzyme +” conditions in HEPES (pH 7.2) and Tris (pH 7.6) buffers, compared to higher glucometer magnitudes of “DNAzyme −” conditions, may indicate a large difference and therefore could appear as a desirable outcome. However, a complete absence of glucometer readings at DNAzyme concentrations as high as nM may cause a misleading outcome in an actual detection scenario involving lower analyte and (by extension) lower DNAzyme concentration. Thus, the combined findings from the glucometer and absorbance highlighted that MES pH 5.0 buffer would support the DNAzyme-catalyzed oxidation of ferrocyanide. At the same time, HEPES (to some extent) and Tris (largely) failed to support the DNAzyme-catalyzed oxidation of ferrocyanide. Most importantly, the parallel assessment of absorbance and glucometer reading underscored the importance of complementary comparative measurements, while establishing that their magnitudes may not always echo each other. Altogether, our outcomes underscored that the ferrocyanide oxidation capability of DNAzyme activity is highly buffer-dependent, even for glucometer readout, and identified 30-40 min duration as the suitable measurement time point.

## Optimization of H_2_O_2_ concentration using absorbance

So far, it was identified that both MES (pH 5.0) and HEPES (to some extent) (pH 7.2) had provided the most suitable conditions for a DNAzyme in catalyzing ferrocyanide to ferricyanide oxidation. They also observed a significant difference both in terms of absorbance and glucometer (more so for MES) readings in the presence and absence of DNAzyme. Having assessed the suitability of the buffer and pH, we next aimed to determine the catalytic ferrocyanide oxidation activity of the DNAzyme under various hydrogen peroxide (H_2_O_2_) concentrations in terms of absorbance under these buffer conditions (Figure 4, S2, and S3). We noted a significant difference in absorbance %fold change at the lower (2.5, 5, and 10 mM) H_2_O_2_ concentrations between the presence/absence of DNAzyme in both MES pH 5.0 and HEPES pH 7.2. However, with increasingly higher H_2_O_2_ concentrations in both MES and HEPES, the %fold change difference between “DNAzyme present” conditions remained almost similar, while those between the “DNAzyme present/absent” consistently decreased (Figure 4B and C). However, the differential change at any H_2_O_2_ concentration in HEPES always remained lower compared to its MES counterpart (Figure 4C). The representative time course plots for MES demonstrated that an increasing starting (0 min) absorbance for “DNAzyme present” condition for lower H_2_O_2_ concentrations (Figure S2 and Figure S3). This was in sync with our hypothesis that DNAzyme rapidly sequestered H_2_O_2_ to generate a “burst” of ferricyanide within probably the first few hundred milliseconds. This also indicated that the reduced difference in %fold change at higher concentrations of H_2_O_2_ could occur due to reduced DNAzyme catalytic efficiency (Figure 4B), which may originate from oxidation of dGs in the G-quadruplex in an increasingly oxidizing environment. Alternatively, this could also be due to competitive loss of substrate for DNAzyme, as higher H_2_O_2_ presence kept on non-catalytically oxidizing the ferrocyanide, reducing the %fold change difference between the presence and absence of DNAzyme. In contrast, similar 0 min absorbance but lesser absorbance %fold change in HEPES buffer suggested lesser DNAzyme catalytic activity under these conditions (Figure 4C and S3). Overall, this firmly established that the MES buffer at lower H_2_O_2_ concentrations (2.5, 5, and 10 mM) provides the most favourable conditions for DNAzyme catalysis of ferrocyanide to ferricyanide.

**Figure 4.**
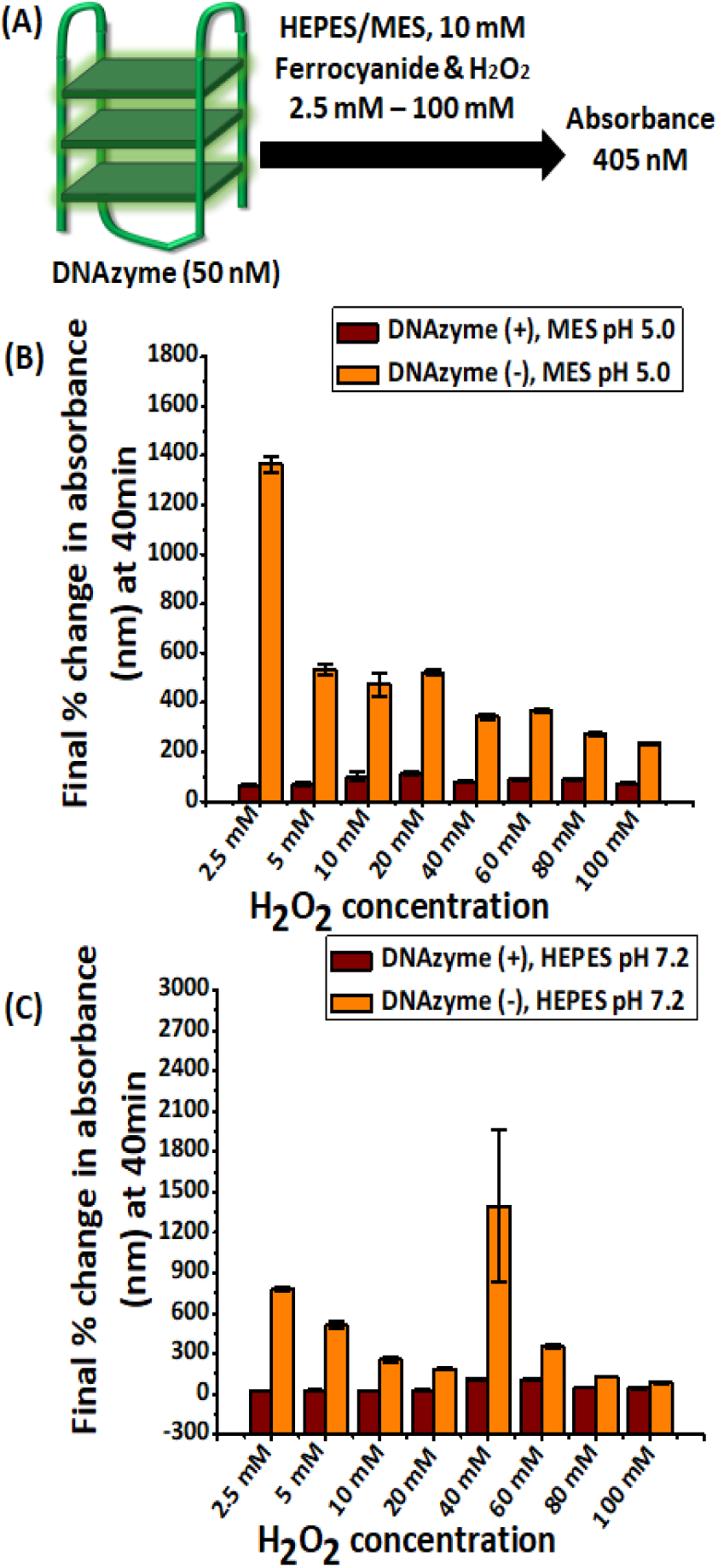
Optimization of H_2_O_2_ concentration (2.5 – 100 mM) for 50 nM DNAzyme-catalysed oxidation of potassium ferrocyanide (10 mM) to ferricyanide. Panel A illustrates the schematic representation of the experimental setup. Panel B and C show the final percentage change in absorbance (405 nm) between 0-40 minutes in different buffer systems: MES pH 5.0 (Panel B) and HEPES pH 7.2 (Panel C). Error bars represent standard deviation (n = 3).

### Investigation of H_2_O_2_ concentration for glucometer readout

Having optimized the suitable H_2_O_2_ concentrations at MES (pH 5) and HEPES (pH 7.2), we investigated whether the differential absorbance measurable for the presence/absence of DNAzyme activity would similarly be reflected in glucometer reading. To achieve this, we tested DNAzyme activity in MES pH 5.0 and HEPES pH 7.2 buffer in 5 mM and 10 mM of H_2_O_2_ concentration in terms of glucometer reading (mg/dL) (Figure 5 and Figure S4). The glucometer reading showed that DNAzyme activity in MES pH 5.0 was detectable by 30 and 20 min for 5 and 10 mM H_2_O_2,_ but never appeared for HEPES (Figure 5B and S4B-D). In addition, the readings reaffirmed that DNAzyme presence/absence from a MES-based assay can be conclusively decided by a 40 min reading. In addition, we experienced reading fluctuations across replicate magnitudes in HEPES (not shown). While 2.5 mM H_2_O_2_ demonstrated higher differential absorbance %fold change for presence/absence of DNAzyme (Figure 4D), neither condition actually provided any readings in the glucometer (data not shown). This was because the 2.5 mM absorbance readings (Figure S2B and S3B), even at 40 min, were less in magnitude than the lowest perceivable glucometer readings (Figure S1). Similarly, reactions conducted at even 5 mM H_2_O_2_ in HEPES buffer did not record in any glucometer readouts for either DNAzyme presence/absence conditions (data not shown). Overall, this implies that MES buffer pH 5.0 is more suitable for supporting DNAzyme catalysis activity in H_2_O_2_ concentrations of 5 mM and 10 mM. Due to observed sub-optimal performance in terms of both absorbance and glucometer usage, the HEPES condition was not pursued further.

**Figure 5.**
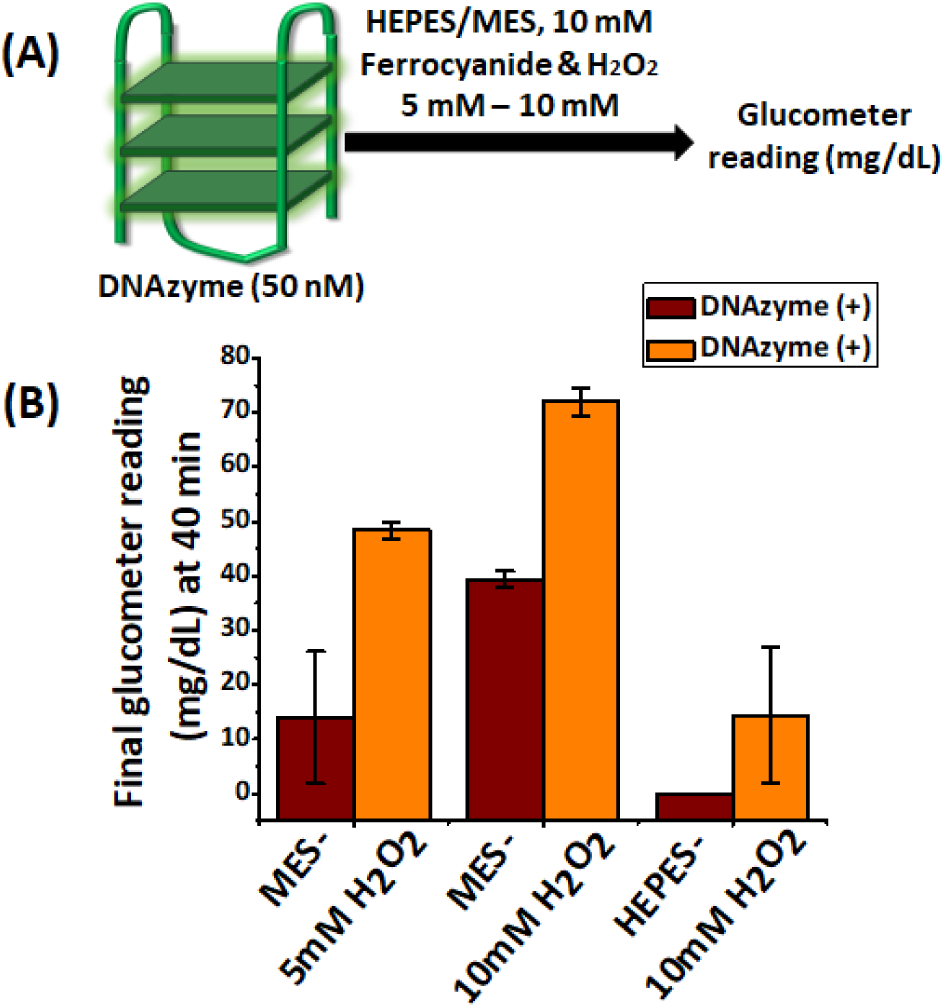
Effect of varying H_2_O_2_ concentration and buffer on DNAzyme-catalyzed oxidation of potassium ferrocyanide to ferricyanide monitored by glucometer. Panel A, Schematic illustration of the experimental setup. Panel B, Final glucometer reading (mg/dL) recorded at 40 minutes under different experimental conditions: 5 mM H_2_O_2_ (MES buffer pH 5.0) or 10 mM of H_2_O_2_ (MES pH 5.0 or HEPES pH 7.2), each containing 10 mM ferrocyanide in the presence and absence of DNAzyme. Error bars represent standard deviation (n = 3). Experiments were performed with Dr. Morepen GLuco One brand.

### Optimization of ferrocyanide concentration using absorbance and glucometer

Next, we probed the optimal ferrocyanide concentration at which the highest differential magnitude for the presence/absence of DNAzyme activity in the now standardized H_2_O_2_ (5 mM and 10 mM) and MES buffer could be obtained. Towards this objective, we experimented with varying concentrations of ferrocyanide (1 – 50 mM) in an optimized MES buffer pH 5.0 system at either 5 mM or 10 mM H_2_O_2_. Once again, our goal was to ascertain the concentration(s) at which the highest absorbance %fold change between the presence/absence of DNAzyme activity at 0-40 min could be obtained (Figure 6). The experiment demonstrated that at 5 mM H_2_O_2_, ferrocyanide at 4, 6, and 8 mM resulted in the highest differential absorbance 0-40 min %fold change for the presence/absence of DNAzyme, with the %fold change difference steadily decreasing with increasing ferrocyanide concentration (Figure 6B and S5). Similarly, at 10 mM of H_2_O_2_, the maximum absorbance %fold change difference between DNAzyme presence/absence was observed at ferrocyanide concentrations of 1, 4, 6, and 8 mM (Figure 6C and S6), after which it steadily decreased. Interestingly, the difference between the presence/absence of DNAzyme in terms of fold change was generally less in 10 mM H_2_O_2_, presumably due to greater non-catalytic oxidation of ferrocyanide.

**Figure 6.**
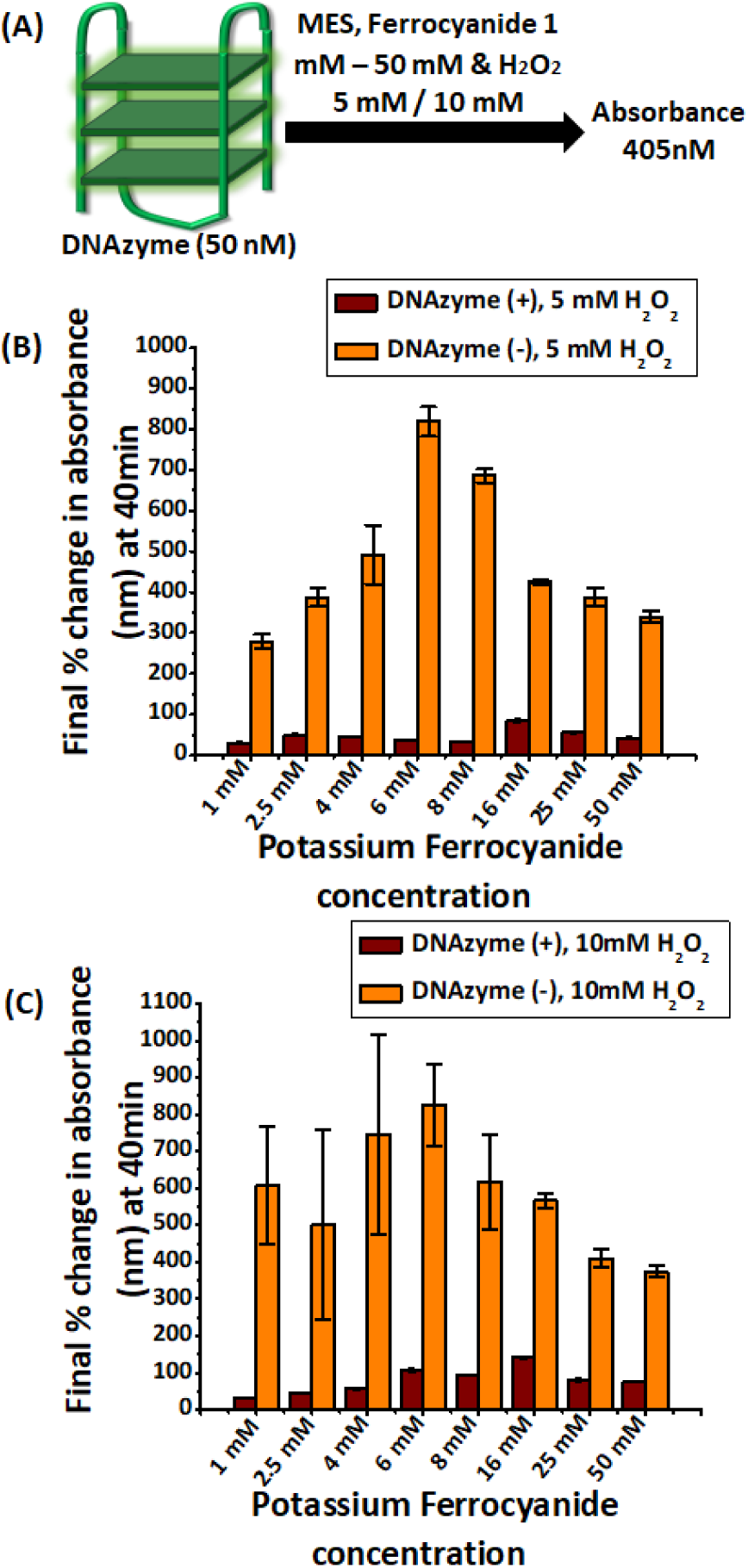
Effect of ferrocyanide concentration on DNAzyme-catalysed oxidation of potassium ferrocyanide to ferricyanide. Panel A, Schematic representation of the experimental setup. Panel B and C, Final percentage change in absorbance (405 nm) measured between 0-40 minutes in MES buffer (pH 5.0). Ferrocyanide oxidation was measured under two optimized H_2_O_2_ concentrations, 5 mM (Panel B) and 10 mM (Panel C), with varying concentrations of ferrocyanide in the presence or absence of 50 nM DNAzyme. Error bars represent standard deviation (n = 3).

Another key observation for “DNAzyme present” conditions in both 5 and 10 mM H_2_O_2_ was reaching an absorbance plateau at the very beginning of the assay for lower ferrocyanide concentrations (generally 1-8 mM) (Figure S5B-D and S6D-F). This was in contrast with an initial slope followed by plateaued absorbance for higher ferrocyanide concentration (16 – 50 mM) (Figure S5E-I, S6G-I). In addition, starting (0 min) absorbances were already significantly higher (0.25 – 0.3) for all ferrocyanide concentrations for both 5 and 10 mM H_2_O_2,_ in contrast to “DNAzyme absence” absorbance (generally less than 0.1) (Figure S5B-I and S6B-I). This presumably suggests that for lower ferrocyanide conditions, DNAzyme would quickly (probably even before the 0 min reading) oxidize ferrocyanide and reduce H_2_O_2_. At this stage, the available H_2_O_2_ or available ferrocyanide or both was not significant enough to generate enough ferricyanide to cause an absorbance change either by catalytic DNAzyme activity or non-catalytic oxidation. At the higher ferrocyanide concentration, the DNAzyme was probably consuming ferrocyanide till 10-15 min before reaching plateaued activity. Overall, this implies that the DNAzyme catalytic efficiency is dependent on ferrocyanide concentrations to achieve maximum activity at each H_2_O_2_ level. Altogether, this suggested a balance is required between ferrocyanide (4, 6, and 8 mM) and H_2_O_2_ concentrations (5 and 10 mM) for achieving an optimal DNAzyme performance.

### Screening DNAzyme operational concentration range in absorbance and glucometer

Having optimized the range of buffer, pH, ferrocyanide, and H_2_O_2_, we next probed the DNAzyme operational concentration range where its presence/absence can be reliably distinguished through absorbance and glucometer readings. By estimating the DNAzyme activity range for these two readouts, these experiments would be expected to serve as guidance to design robust DNAzyme transducer-based bioanalytical assays. To achieve this objective, we conducted an experiment using standardized ferrocyanide concentrations (4, 6, and 8 mM) with a fixed H_2_O_2_ concentration (5 or 10 mM). DNAzyme concentrations ranging from 1 pM - 200 nM were initially tested to evaluate their effect on the absorbance %fold change at 0-40 minutes (Figures 7, 8, S7-S12). Overall, the experiments demonstrated that increasingly smaller differences for 0-40 min absorbance %fold change between DNAzyme presence/absence conditions with successively lowered DNAzyme concentrations. Moreover, the same DNAzyme concentration had increasingly greater absorbance %fold change difference from “DNAzyme absent” conditions with higher ferrocyanide concentration. For 1 pM DNAzyme concentration at 4, 6, 8 mM ferrocyanide and 5 mM H_2_O_2_ concentration, for instance, the %fold change difference between DNAzyme presence/absence was 2.95x, 3.33x, and 5x (Figure 7B-C). Generally similar observations about absorbance %fold change can be noted for DNAzyme presence/absence conditions in 10 mM H_2_O_2_ as well (Figure 8). However, unlike 5 mM H_2_O_2_, the absorbance %fold change for 1-10 pM DNAzyme presence/absence was undifferentiable in 10 mM H_2_O_2_ (Figure 8), possibly due to non-catalytic H_2_O_2_-mediated oxidation of ferrocyanide.

**Figure 7.**
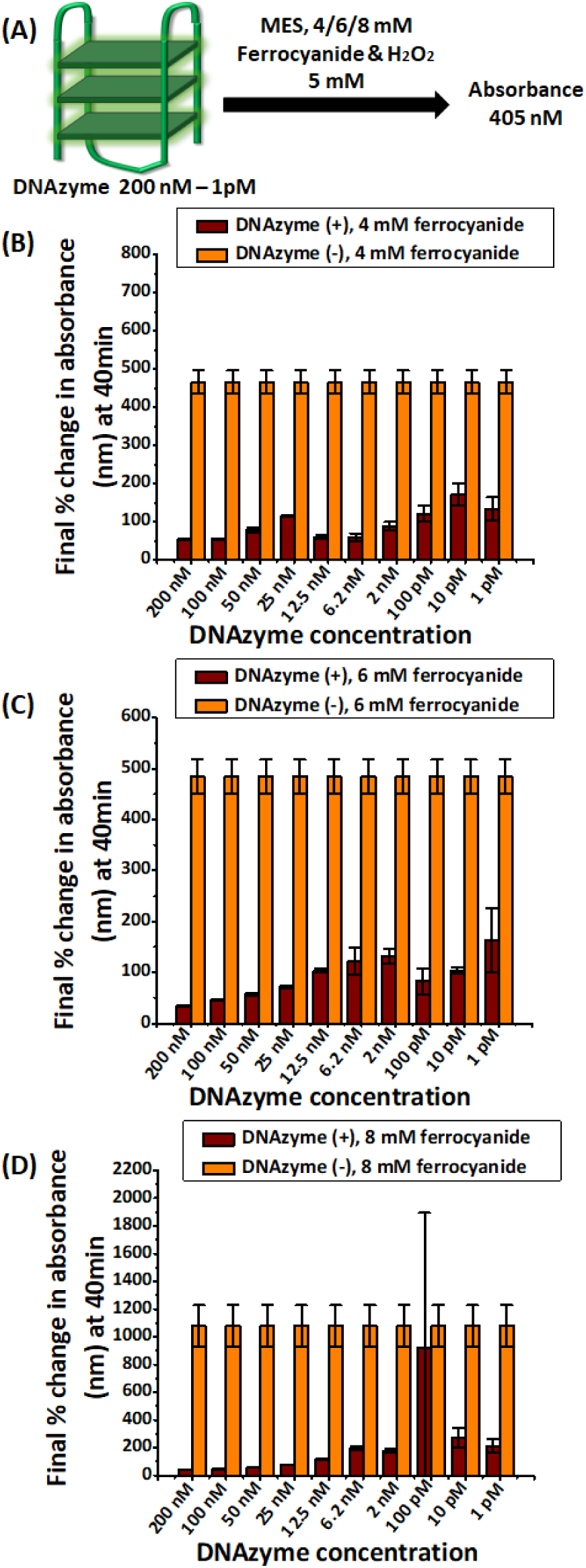
Optimization of DNAzyme concentration for DNAzyme-catalyzed oxidation of potassium ferrocyanide to ferricyanide. Panel A provides a schematic representation of the experimental setup. Panel B-D shows the %fold change in absorbance (405 nm) measured at 0-40 minutes in MES buffer (pH 5.0) with a constant 5 mM concentration of H_2_O_2_ and ferrocyanide concentrations of 4 mM (Panel B), 6 mM (Panel C) and 8 mM (Panel D) with and without DNAzyme. As ferrocyanide and H_2_O_2_ conditions were same in each individual panel, the magnitudes for absorbance %fold change for “DNAzyme−” condition were kept identical during plotting for individual Figure panels (but separate across panels). Error bars represent standard deviation (n = 3).

**Figure 8.**
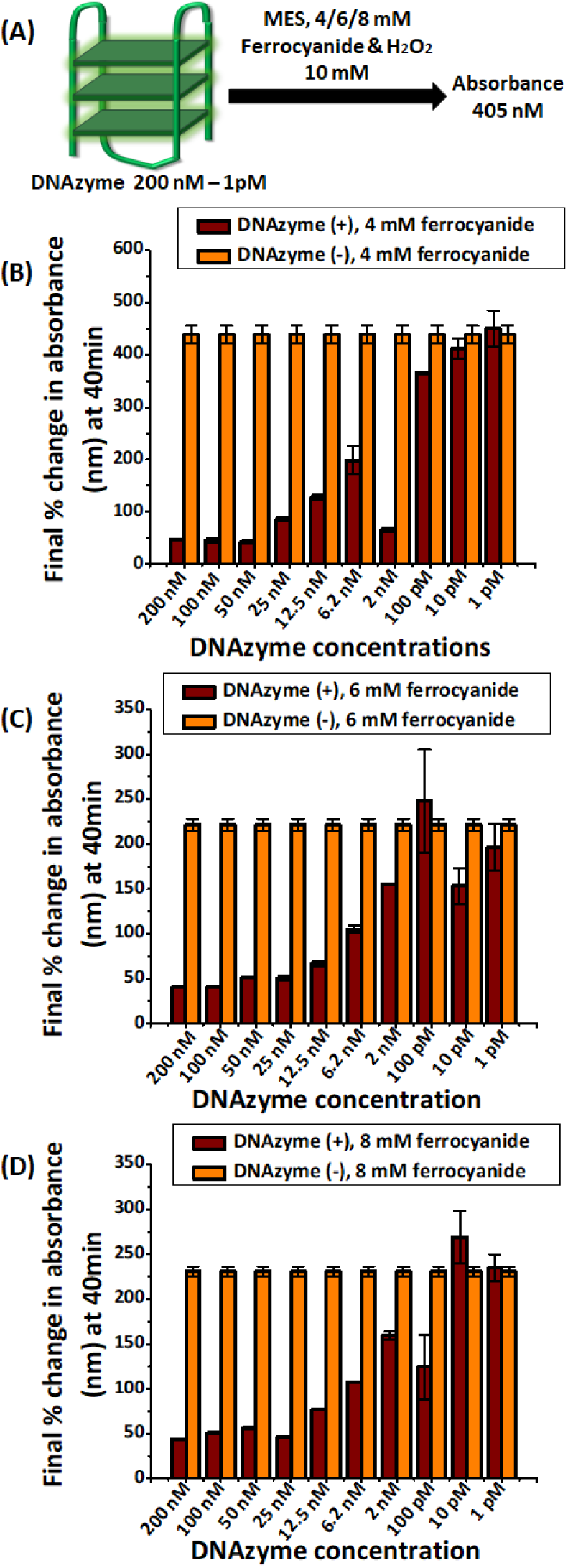
Optimization of DNAzyme concentration range for DNAzyme-catalyzed oxidation of potassium ferrocyanide to ferricyanide. Panel A, Schematic representation of the experimental setup. Panel B-D, %fold change in absorbance measured at 0-40 minutes in MES buffer (pH 5.0) with a constant 10 mM concentration of H_2_O_2_ and ferrocyanide concentrations of 4 mM (Panel B), 6 mM (Panel C) and 8 mM (Panel D) with and without DNAzyme (1 pM – 200 nM). As ferrocyanide and H_2_O_2_ conditions were same in each individual panel, the magnitudes for absorbance %fold change for “DNAzyme−” condition were kept identical during plotting for individual Figure panels (but separate across panels). Error bars represent standard deviation (n = 3).

An interesting observation from the time course of absorbance studies was that the initial (0 min) absorbance for higher DNAzyme concentrations (12.5 – 200 nM) was greater (0.08 to 1) than respective “DNAzyme absent” conditions (less than 0.05) (Figure S7B-F). After an initial increment upto 10-15 min, the absorbance for “DNAzyme present” plateaued. For lesser DNAzyme concentrations (1 pM – 6.2 nM), the starting (0 min) absorbance was either comparable or less than the respective “DNAzyme absent” conditions (Figure S7G-K). In our opinion, a higher amount of DNAzyme (12.5 – 200 nM) oxidized a significant amount of ferrocyanide to generate ferricyanide even before the 0 min observation point, causing large initial absorbance. The later slowdown was either due to loss of catalytic activity or absence of an ample amount of substrate (ferrocyanide or H_2_O_2_ or both). In comparison, a lower amount of DNAzyme continued to consume ferrocyanide throughout the assay time, thus having low absorbance throughout the assay but still generating higher absorbance %fold change than higher DNAzyme concentrations (Figures 7B-D, S7H-I, S8H-I, S9H-I). A similar trend was observable for low DNAzyme concentrations in 4-8 mM ferrocyanide in 10 mM H_2_O_2_ as well (Figures S10G-K, S11H-K, S12H-K). Under such conditions, low absorbance magnitudes but greater 0-40 min %fold change were discernible. Overall, the %fold change in absorbance at 0-40 minutes provided critical insight into the extent of ferrocyanide to ferricyanide formation in the presence and absence of DNAzyme. It also provided a glance into a possible working range of the hemin DNAzyme as a transducer under these conditions.

Having identified a possible working range for the DNAzyme transducer through absorbance measurement, we next determined whether this range would translate to glucometer readout for effective differentiation between DNAzyme presence/absence conditions. Towards this objective, we conducted the same experiment using the already standardized ferrocyanide concentrations (4, 6, and 8 mM) with a fixed H_2_O_2_ concentration (5 or 10 mM) (Figures 9 and 10). As optimized earlier, 1 pM – 200 nM DNAzyme concentrations were evaluated on the glucometer reading at 40 min for DNAzyme presence/absence. For all three ferrocyanide concentrations, lowering DNAzyme concentration from 200 nM first led to a gradual lowering of the difference with its “DNAzyme absent” counterpart to various degrees. In case of 4 nM ferrocyanide in 5 mM H_2_O_2_, for instance, this difference persisted till 25 nM, while for 6 and 8 mM, it was till 100 and 50 nM (Figure 9B-D and Figure 10B-D).

**Figure 9.**
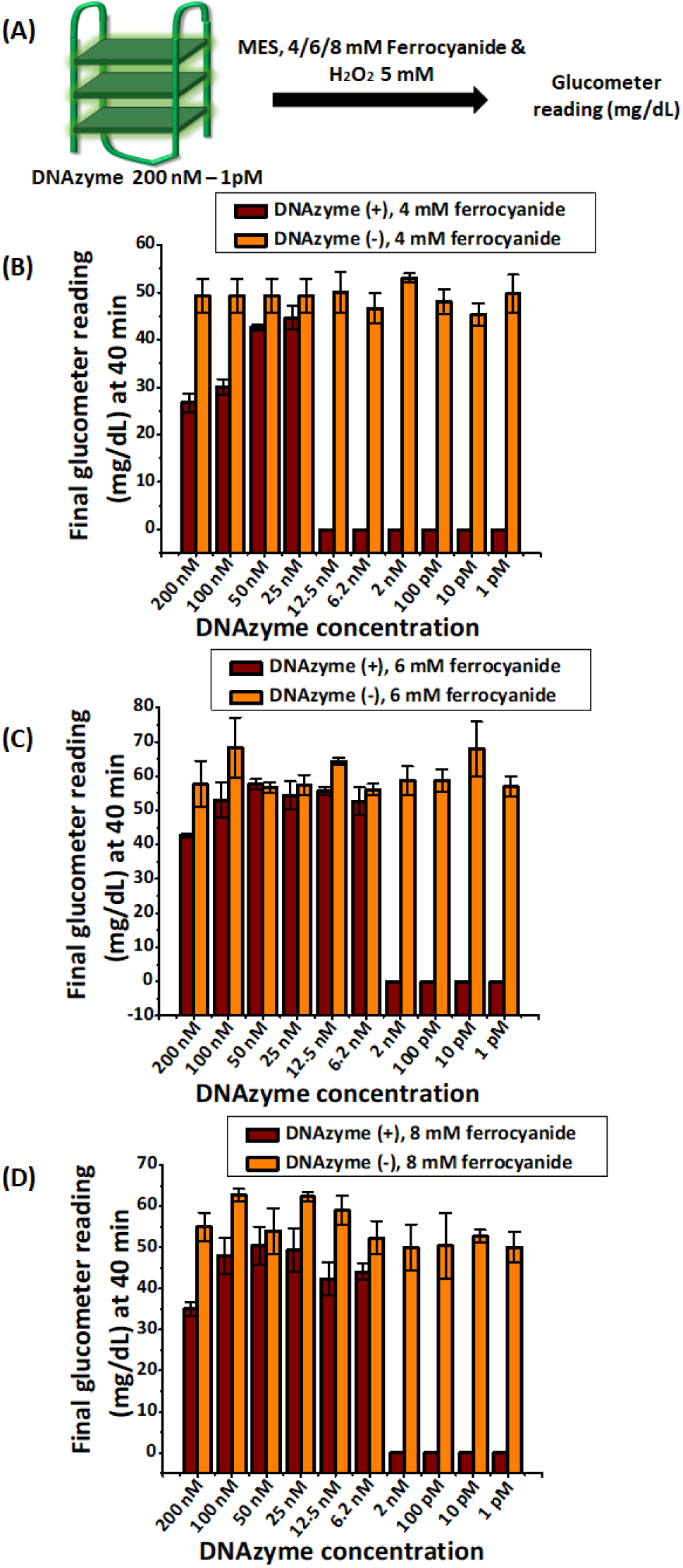
Optimization of DNAzyme concentration range for DNAzyme-catalyzed oxidation of potassium ferrocyanide to ferricyanide using glucometer. Panel A provides a schematic representation of the experimental setup. Panel B-D shows the final glucometer reading (mg/dL) at 40 minutes in MES buffer (pH 5.0) with a constant 5 mM concentration of H_2_O_2_ and ferrocyanide concentrations of 4 mM (Panel B), 6 mM (Panel C) and 8 mM (Panel D) with or without DNAzyme (1 pM – 200 nM). Error bars represent standard deviation (n = 3). Experiments were performed with Dr. Morepen GLuco One brand.

**Figure 10.**
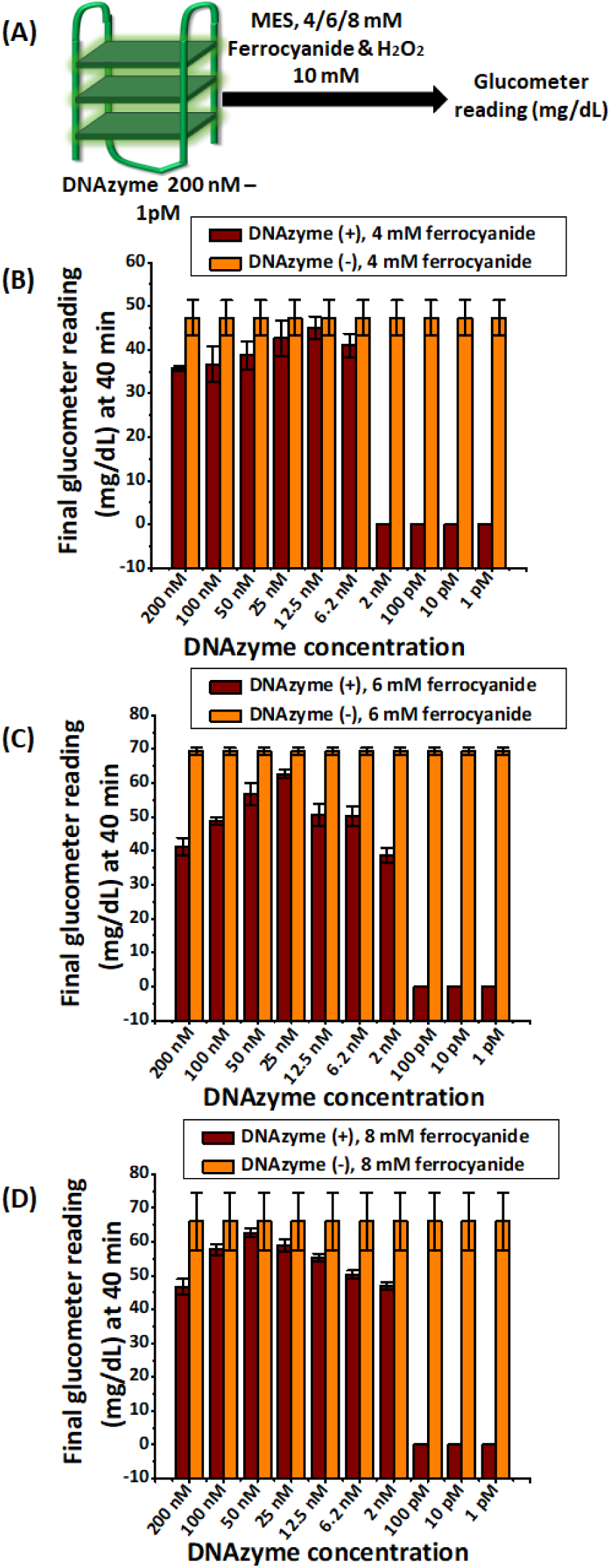
Optimization of DNAzyme concentration range for DNAzyme-catalyzed oxidation of potassium ferrocyanide to ferricyanide using glucometer. Panel A, Schematic representation of the experimental setup. Panel B-D shows the final glucometer reading (mg/dL) at 40 minutes in MES buffer (pH 5.0) with a constant 10 mM concentration of H_2_O_2_ and ferrocyanide concentrations of 4 mM (Panel B), 6 mM (Panel C) and 8 mM (Panel D) with or without DNAzyme (1 pM – 200 nM). As ferrocyanide and H_2_O_2_ conditions were same in each individual panel, the magnitudes for glucometer reading for “DNAzyme−” condition were kept identical during plotting for individual Figure panels (but separate across panels). Error bars represent standard deviation (n = 3). Experiments were performed with Dr. Morepen GLuco One brand.

Interestingly, the difference between DNAzyme present/absent conditions for further lower concentrations (for e.g., 1 pM - 12.5 nM DNAzyme for 4 mM ferrocyanide) became much significant for all three ferrocyanide conditions (Figure 9B-D and Figure 10B-D). Although self-contradictory and perplexing, these observations could be rationalized from the absorbance studies (Figures 7B-D, 8B-D, and S7-S12). In case of 4 mM ferrocyanide in 5 mM H_2_O_2_, for example, the 0-40 min absorbance %fold change was higher for lower DNAzyme concentrations (for instance, at 1 pM) compared to its higher “DNAzyme present” (e.g., 200 nM) counterpart. But at the same time, it was lower than “DNAzyme absent” conditions (Figure 7B, Figure S7B and S7K). As explained in the previous section, this would indicate continued DNAzyme activity for the measurement duration (0-40 min) for low (1 pM) DNAzyme concentration. However, despite continued DNAzyme activity, it still did not generate enough ferricyanide for a glucometer reading. The 40 min absorbance reading for 1 pM DNAzyme, for instance, was below 0.02 (see Figure S7K). As evident from the absorbance vs glucometer reading vs concentration standard curve at Figure S1, this was insufficient to generate a glucometer reading. Therefore, the DNAzyme transducer at such a low concentration would act as a qualitative yes/no with a glucometer readout. For an actual sensing application involving the hemin DNAzyme as a transducer and a glucometer as a readout, the absence of a glucometer readout at 40 min would therefore indicate the presence of the analyte.

### Screening DNAzyme operational concentration range below pM concentration

Intrigued by a qualitative albeit more sensitive at lower DNAzyme concentration, we explored the lowest possible concentration at which DNAzyme transducer activity would confer a differential readout for a glucometer. To pursue this objective, we explored the time course (0-40 min) absorbance change for DNAzyme presence/absence and glucometer readings for 1 aM – 100 fM DNAzyme concentrations in 4 mM ferricyanide and 5 mM H_2_O_2_. Interestingly, and similar to 1-10 pM DNAzyme concentrations, glucometer readings were again successful in differentiating DNAzyme presence/absence till aM (Figure 11C). At the same time and perhaps expectedly, 0-40 min absorbance %fold change was undifferentiable (Figure 11B). Once again, this perplexing finding was rationalizable from the absorbance time course plots (Figure S13B-H). While the “0 min” absorbance magnitude for “DNAzyme present” condition was lower than the respective “DNAzyme absent”, the differential absorbance growth curve indicated continued DNAzyme activity till at least 10-100 aM concentration. Possibly the initial burst of activity (to consume ferrocyanide), followed by low but persistent DNAzyme activity over 40 min experiment duration, failed to generate enough ferricyanide (less than even a non-catalytic “DNAzyme absent” control) to elicit a glucometer response. Altogether, this study established the optimal operational DNAzyme concentration range for transducer activities.

**Figure 11.**
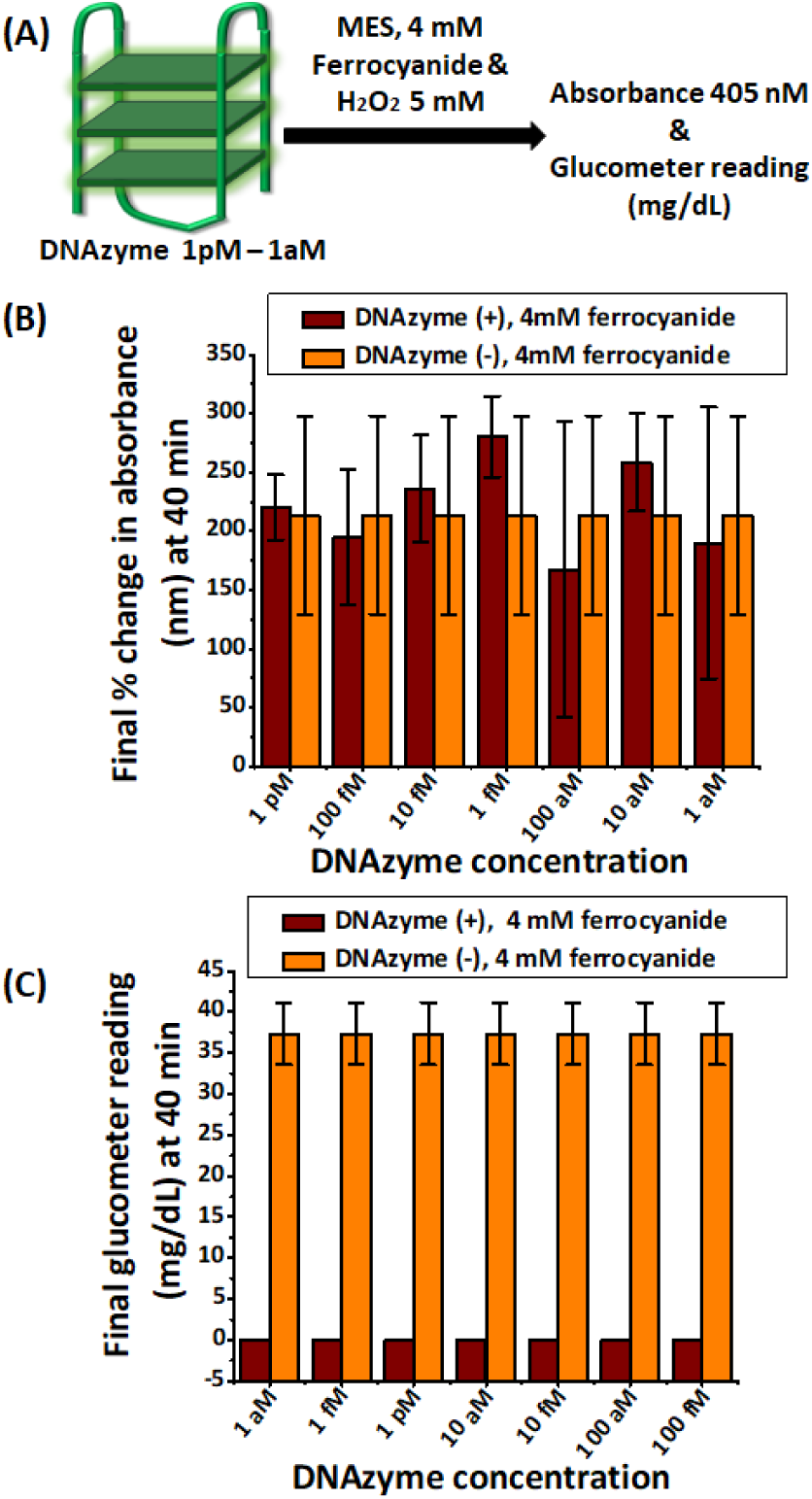
Optimization of DNAzyme concentration range for DNAzyme-catalyzed oxidation of potassium ferrocyanide to ferricyanide. Panel A, Schematic representation of the experimental setup. Panel B, %Fold change in absorbance measured at 0-40 minutes in MES buffer (pH 5.0), with a constant 5 mM concentration of H_2_O_2_ and ferrocyanide at 4 mM. Panel C, Final glucometer reading (mg/dL) at 40 minutes in MES buffer (pH 5.0) with a constant 5 mM concentration of H_2_O_2_ and ferrocyanide concentration of 4 mM with or without DNAzyme (1 aM – 100 fM). As ferrocyanide and H_2_O_2_ conditions were same in each individual panel, the magnitudes for absorbance %fold change and glucometer for “DNAzyme−” condition were kept identical during plotting for individual Figure panels (but separate across panels). Error bars represent standard deviation (n = 3). Experiments were performed with Dr. Morepen GLuco One brand.

### Observation of DNAzyme transducer activity in an alternative glucometer brand and use of HRP as a transducer for glucometers

We wanted to verify if this mechanism would be translatable in alternate glucometer brands as well, and whether there would be a significant difference in readout. To investigate this, we tested DNAzyme activity (concentrations 1 pM – 1 aM) using the glucometer brand “Apollo pharmacy blood glucose monitoring” with 4 mM ferrocyanide and 5 mM H_2_O_2_, while readings were taken at 40 min. Before proceeding, we first confirmed that this particular brand could also detect and differentiate variable ferricyanide concentrations as a “glucose” readout in mg/dL units (data not shown). Similar to “Dr. Morepen GLuco One brand” brand from the above studies, the “Apollo pharmacy blood glucose monitoring” brand also demonstrated differential measurement between DNAzyme present/absent conditions till aM range (Figure 12). This experiment thus established that the assay conditions are “transferrable” across other commercial glucometer brands that are responsive to ferricyanide measurements.

**Figure 12.**
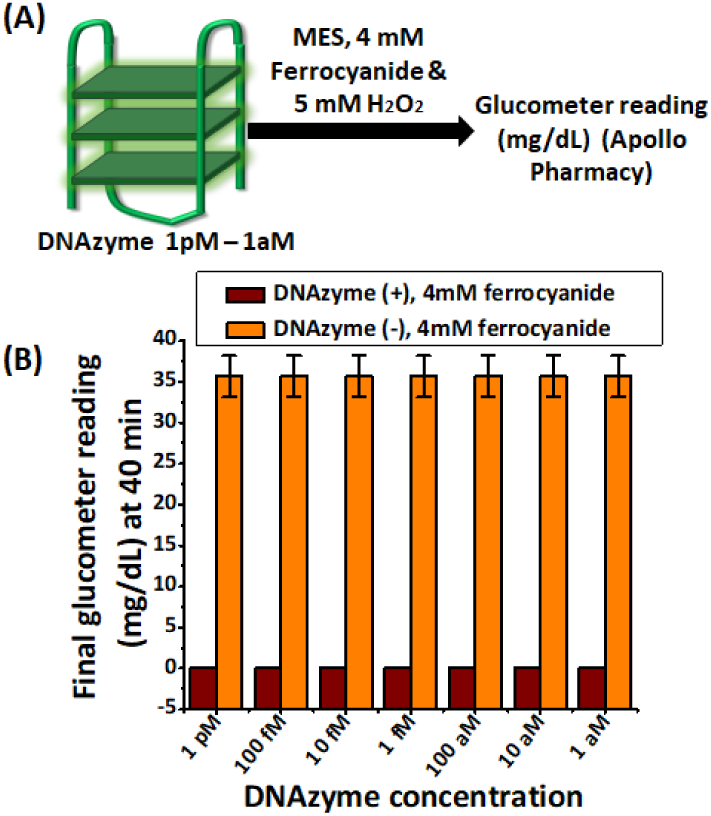
Optimization of DNAzyme concentration range for DNAzyme-catalyzed oxidation of potassium ferrocyanide to ferricyanide using “Apollo pharmacy” glucometer. Panel A, Schematic representation of the experimental setup. Panel B, Final glucometer reading (mg/dL) at 40 minutes in MES buffer (pH 5.0) with a constant 5 mM concentration of H_2_O_2_ and ferrocyanide concentrations of 4 mM with or without DNAzyme concentration ranging from 1 aM to1 pM. As ferrocyanide and H_2_O_2_ conditions were same in each individual panel, the magnitudes for glucometer reading for “DNAzyme−” condition were kept identical during plotting for individual Figure panels (but separate across panels). Error bars represent standard deviation (n = 3). Experiments were performed with “Apollo pharmacy blood glucose monitoring” glucometer.

Next, we explored whether HRP activity would be detectable in “Dr. Morepen GLuco One brand” glucometer with ferrocyanide as a substrate. If successful, this proof-of-concept study would open up limitless possibilities to explore the direct detection of even ELISA readouts via a glucometer. To decide a possible HRP concentration range, we investigated ABTS substrate-mediated chromogenic activity for HRP concentrations 1 nM – 10 μM (Figure 13A). Barring 1 nM, all other concentrations generated a characteristic dark green color within 2-5 min. These HRP concentrations were then subjected to glucometer measurement in experimental conditions optimized for DNAzyme activity measurement (4 mM ferrocyanide, 5 mM H_2_O_2_, MES pH 5.0, along with Na and K salts). The glucometer readings taken at 0, 10, and 30 min were all able to differentiate between HRP present/absent conditions (Figure 13B-E). While we are currently engaged in investigating precise experimental conditions for optimal HRP activity parameters, this preliminary study definitively indicates that HRP activity (and by extension, ELISA) would be measurable through a glucometer using ferrocyanide as redox substrate.

**Figure 13:**
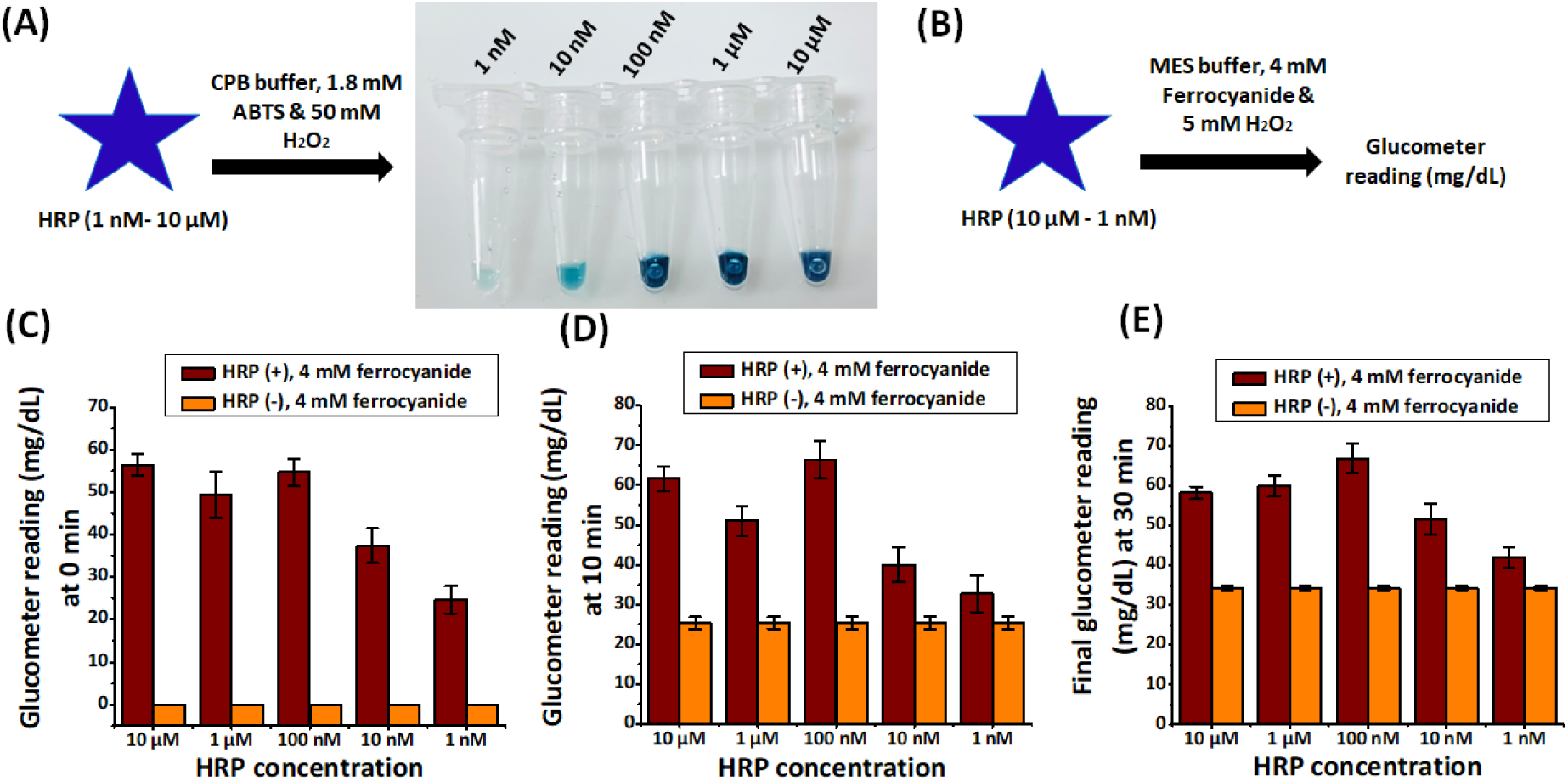
Colorimetric and oxidation activity of HRP activity at varying enzyme concentrations. Panel A, Colorimetric analysis of HRP activity at concentrations 1 nM to 10 µM in CPB pH 4.5, in the presence of 1.8 mM ABTS (2,2’-azino-bis(3-ethylbenzothiazoline-6-sulfonic acid)) and 50 mM hydrogen peroxide (H₂O₂). Panel B, Schematic representation of oxidation activity of HRP (1 nM – 10 μM) monitored by glucometer readings. Panel C-E, Catalytic oxidation of ferrocyanide measured with glucometer with or without HRP (1 nM to 10 µM) in MES buffer pH 5.0 containing a 4 mM ferrocyanide, 5 mM H_2_O_2_. Glucometer readings were taken at 0 (Panel C),10 (Panel D), or 30 min (Panel E). Error bars represent standard deviation (n = 3). Experiments were performed with Dr. Morepen GLuco One brand.

## Discussion

This study investigates the hypothesis whether an over-the-counter glucometer could be used as a general-purpose readout instrument for assays that involve HRP analogues such as hemin DNAzyme. We aimed to do so without any chemical conjugation between the analogues and biomolecular recognition elements such as antibodies or aptamers. Furthermore, to keep the assay user-friendly, low-cost, and widely adoptable, we also wanted to minimize the use of unusual and expensive redox intermediaries. The hypothesis is founded on the basis of early literature documenting potassium ferrocyanide as a redox substrate of HRP and widespread utilization of ferrocyanide/ferricyanide couple in commercial glucometer strips^20,23^. Towards this objective, we used a commonly utilized analogue of HRP, HRP-mimic hemin DNAzyme, as our biocatalyst and probed the role of buffer, pH, H_2_O_2_, and redox mediator (ferrocyanide) concentration. To closely simulate actual ELISA/ELASA-like assays and to remove excess hemin, we immobilized the DNAzyme on streptavidin magnetic beads. Furthermore, the reaction kinetics were studied through absorption spectral monitoring of potassium ferricyanide (the product in this assay) generation to gain an in-depth understanding of the optimization of each parameter. Hemin DNAzymes, either by themselves, in conjugation with nucleic acid binder, or through being embedded in various nanoparticles, have been extensively utilized for detecting nucleic acid amplification, aptameric biorecognition, metal ions, and small molecules^25–29^. However, these applications were limited to DNAzyme use as chromogenic, fluorogenic, or electrogenic transducers involving centralized instruments such as multimode plate readers. Therefore, establishing a glucometer-based readout for hemin DNAzyme transducer would facilitate a decentralized, low-cost, and extremely user-friendly tool towards diverse sensing applications.

Our study unequivocally establishes that HRP-mimic hemin DNAzyme, in the presence of H_2_O_2_, can oxidize ferrocyanide to ferricyanide. The latter, being a measurable substrate of a conventional glucometer via its strips, enables the detection of DNAzyme activity in a glucometer. We investigated the feasibility of the reaction in MES (pH 5.0), sodium acetate (pH 5.0), HEPES (pH 7.2 and 8.2), and Tris buffer (pH 7.6 and 8.5) through both ferricyanide generation (measured using absorption spectroscopy) and glucometer readout. As the MES condition seemed most suitable for both these measurements, it was selected for carrying out further studies for H_2_O_2_ and ferrocyanide concentration optimization. Using optimized buffer, pH, H_2_O_2_, and ferrocyanide concentration, we then demonstrated that the aforementioned DNAzyme activity is measurable in a glucometer between sub-micromolar to aM concentration range. Additionally, the activity could be detected in at least two commercial glucometer brands, providing equivalent readouts. Finally, our preliminary experiments involving HRP suggest that the optimized experimental conditions could be extended towards successful glucometer-based HRP activity measurement as well. Altogether, our study investigated the modalities of using HRP-mimic hemin DNAzyme as a transducer for ELISA or ELASA-like assays.

Use of glucometer in ELISA, ELASA, or related assays involving either aptamer or antibodies as biorecognition elements has utilized invertase or glucoamylase as the transducer^21,22^. Usually chemically conjugated to the secondary antibody or the nucleic acid recognition domain, these enzymes generate glucose, which is measurable in a glucometer. Several subsequent studies have extensively bioengineered this assay to detect several diverse biomolecules, small molecules, as well as molecular pathways^33^. In addition, these enzymes, in conjunction with antibodies or nucleic acid probes, have been utilized as genosensors or reporters for nucleic acid amplification as well^34,35^. Our assay bypassed the glucose generation pathway and instead explored the direct utilization of ferrocyanide (a redox mediator measurable in a glucometer) by hemin DNAzyme as a substrate. Although ferrocyanide was identified as a substrate for HRP in earlier studies^23^, those studies predominantly focused on understanding the redox intermediates of HRP itself using stopped flow kinetics. Our study (for the first time, to the best of our knowledge) demonstrated that HRP-mimic hemin DNAzyme could also utilize ferrocyanide as redox mediator in the presence of H_2_O_2_. Additionally, we (for the first time) also investigated the role of each of the components in this assay and adjusted their concentration accordingly to extract optimal performance. Another very recent study has utilized hemin DNAzyme coupled with Cas12a for detecting nucleic acid presence via glucometer with remarkable analytical sensitivity^36^. In contrast to our study, the aforementioned study, however, does not utilize ferrocyanide directly. It utilized costlier NADH/NAD^+^ redox intermediaries (instead of inexpensive potassium ferrocyanide) as additives that in turn interact with ferrocyanide/ferricyanide redox couple present in glucometer strip to facilitate electron transfer to glucometer strip. Furthermore, unlike our study, the study did not utilize absorption spectroscopy to rationalize reaction dynamics nor probe the role of buffer, pH, concentrations of either H_2_O_2_ or ferrocyanide. As demonstrated in our study, each of these components would play significant role in optimizing the assay condition. The optimization of these parameters are integral and unique to our study.

The use of HRP-mimic hemin DNAzyme has been extensively utilized as transducers of ELISA-like assays for detecting small molecules, ions, nucleic acids, proteins, and whole cell pathogens. This includes Hepatitis B surface antigen^25^, mercury ion^26^, KRAS G12D mutation^27^, *E. coli* O157:H7^28^, micro-RNA^29^ etc. In addition, multimeric hemin DNAzyme transducer motif embedded in a DNA nanostructure for the detection of lead ions^7^, nanoparticle-displaying hemin DNAzyme for the detection of *Salmonella*^37^, surface-immobilized hemin DNAzymes for biosensing of human leptin^38^, DNA nanowire-mounted hemin DNAzyme for prion protein detection^39^, and readouts for isothermal as well as PCR nucleic acid amplification have shown its broad and translational utility^34,35^. While our study extensively catalogues the experimental parametric scope of HRP-mimic DNAzyme as well as a representative HRP usage in glucometer reporter assays, we did not demonstrate its utilization in an actual biosensing assay. This remains one of the limitations of our study. In addition to “Dr. Morepen GLuco One” and “Apollo pharmacy blood glucose monitoring” brands, we have also explored this assay’s utility in another commercial glucometer brand. However, the latter was non-responsive to potassium ferricyanide and (by extension) our assay. We are currently exploring methods on bypassing this limitation to make this assay truly “universal” for all glucometer brands.

As conventional ELISA provides a quantitative readout using linear regression, it is extensively used for highly accurate analyte concentration measurement in biopharma industries. Similarly, identifying a correlation between biomolecule presence and diseases (e.g., lipid levels and the chance of atherosclerosis) requires low-error estimation of biomolecular concentration using ELISA. The technological needs in these scenarios would not be addressable by our innovation. However, semi-quantitative or qualitative disease detection, such as a yes/no presence of a pathogen or even semi-accurate estimation of pathogen load using ELISA, can be determined by our method, contributing to disease detection in remote settings.

We have shown preliminary proof-of-concept evidence that the optimized condition could apply to HRP, and by natural extension, ELISA measurement as well. This prolepsis is founded on the fact that ferrocyanide has been spectroscopically established as an HRP substrate, and its utilization as a substrate was critical in early understanding of the redox behavior of HRP. In addition, a study by Ugarova et al in 1981 demonstrated that utilization of ferrocyanide as a substrate by HRP leads to a relatively modest change in absorbance, while the absence of HRP leads to a faster increase^40^. Another recent study by Lee et al, aiming towards HRP-enabled H_2_O_2_ determination using a glucometer, also utilized ferrocyanide as a substrate^41^. Similar to our study, it demonstrated that peroxide presence caused a relatively lesser magnitude of glucometer reading compared to that for HRP absence. However, it also did not explore the role of pH, buffer, H_2_O_2_, or ferrocyanide/ferricyanide, and the measurements were carried out in Tris pH 7.2 buffer. Our data, as well as these studies, would suggest that HRP, and by extension HRP-based ELISA could as well be utilized for glucometer-enabled readout. Likewise, we anticipate that the activities of various HRP-mimic nanoparticles might also be similarly measurable in glucometer using conditions established in our study. Beyond this, we expect the detection signal could be further boosted by use of ascorbate in the reaction mixture. This is due to fact that that ascorbate can recycle Fe(III) to Fe(II), and therefore ferricyanide (Fe(III)) to ferrocyanide (Fe(II)), thus reinstating the key redox reaction and enhancing the signal intensity further. However for the same reason, it may also cause false positive signal enhancement. Although these possibilities highlight some limitations of our study, they highlight prospective exciting developments concerning glucometer usage in ELISA or similar reactions. Therefore, these are currently being investigated in our lab and will be part of future efforts to explore the full spectrum and depth of these promising directions.

Overall, our study identified that HRP-mimic hemin DNAzyme activity could reliably be measured using a glucometer in a label-free manner and as a robust alternative to ELISA reader, multimode reader, or potentiostats. Utilizing absorbance spectroscopy and glucometer reading, we cataloged buffer, pH, redox substrate, and H_2_O_2_ conditions that would help identify optimal conditions for this assay. Given the widespread availability of glucometers and low cost, it is anticipated that this study will spur future developments towards the use of glucometers for ELISA and ELISA-like assays, leading to democratized access to healthcare.

## Authorship Credits

SP, SG, AT, and MS conceived the experiments and analyses. SP performed the experiments. MS helped understood the enzymology. AT helped understand the molecular diagnosis aspect of the work. SP, SG performed the data analyses. SP and SG wrote the manuscripts. All authors read and approved the manuscript.

## Acknowledgements

This work was supported by the IIIT-iHub grant (No H4-006), DST INSPIRE PhD Fellowship to Ms Saba Parveen (No DST/INSPIRE/03/2021/002361), and Mahindra University internal funding to “Interdisciplinary Centre for Nanosensors and Nanomedicine”. The authors are grateful to Prof. Rajinder S. Chauhan, Prof. Bishnu Pal, and Prof Yajulu Medury (Vice-Chancellor of Mahindra University) for their encouragement and constant scholarly support.

## Supplementary Information

Materials and methods with details experimental protocols; Additional Figures;

