## Supplementary Material for "Evaluating horseradish peroxidase-mimic DNAzyme transducer for glucometer readout: roles of various components and their optimization"

### Table of Content


| Topic | Page No |
|---|---|
| **Materials and Methods** | **S3** |
| **Figure S1.** Correlation between ferricyanide concentration and glucometer readings in MES buffer | **S7** |
| **Figure S2.** Optimization of $H_2O_2$ concentration for DNAzyme-catalyzed oxidation of potassium ferrocyanide in MES | **S8** |
| **Figure S3.** Optimization of $H_2O_2$ concentration for DNAzyme-catalyzed oxidation of potassium ferrocyanide *(10 mM)* to ferricyanide in HEPES | **S9** |
| **Figure S4.** Effect of varying $H_2O_2$ concentration buffer on DNAzyme-catalyzed oxidation of potassium ferrocyanide *(10 mM)* by glucometer. | **S10** |
| **Figure S5.** Optimization of ferrocyanide concentration on DNAzyme-catalysed oxidation of potassium ferrocyanide to ferricyanide *in 5 mM* $H_2O_2$ | **S11** |
| **Figure S6.** Optimization of ferrocyanide concentration on DNAzyme-catalysed oxidation of potassium ferrocyanide to ferricyanide *in 10 mM* $H_2O_2$. | **S12** |
| **Figure S7.** Optimization of DNAzyme concentration *range* for DNAzyme-catalyzed oxidation of potassium ferrocyanide to ferricyanide *(4 mM) in 5 mM* $H_2O_2$ | **S13** |
| **Figure S8.** Optimization of DNAzyme concentration *range* for DNAzyme-catalyzed oxidation of potassium ferrocyanide to ferricyanide *(6 mM) in 5 mM* $H_2O_2$. | **S14** |
| **Figure S9.** Optimization of DNAzyme concentration *range* for DNAzyme-catalyzed oxidation of potassium ferrocyanide to ferricyanide *(8 mM) in 5 mM* $H_2O_2$. | **S15** |
| **Figure S10.** Optimization of DNAzyme concentration *range* for DNAzyme-catalyzed oxidation of potassium ferrocyanide to ferricyanide *(4 mM) in 10 mM* $H_2O_2$. | **S16** |
| **Figure S11.** Optimization of DNAzyme concentration *range* for DNAzyme-catalyzed oxidation of potassium ferrocyanide *(6 mM)* to ferricyanide *in 10 mM* $H_2O_2$. | **S17** |
| **Figure S12.** Optimization of DNAzyme concentration *range* for DNAzyme-catalyzed oxidation of potassium ferrocyanide *(8 mM)* to ferricyanide *in 10 mM* $H_2O_2$. | **S18** |
| **Figure S13.** Optimization of lower DNAzyme concentration *range* for DNAzyme-catalyzed oxidation of potassium ferrocyanide *(4 mM)* to ferricyanide *in* $H_2O_2$. | **S19** |
| **References** | **S19** |

### Materials and Methods

**Materials:**

Potassium ferrocyanide ($K_4[Fe(CN)_6].3H_2O$) (cat # 32294), potassium ferricyanide ($K_3[Fe(CN)_6]$) (cat # 59558), potassium chloride (KCl) (cat # 84984), sodium chloride (NaCl) (cat # 33205), Dimethyl sulphoxide (DMSO) (cat # 24075), triton X-100 (cat # 64518), hemin (cat # 98457), MES monohydrate buffer (cat # 86751) , HEPES Buffer (cat # 16826), sodium acetate (cat # 88035), ABTS (2,2'-Azino-bis(3-ethylbenzothiazoline-6-sulfonic acid) diammonium salt (cat # 40157) were purchased from SRL. Tris (hydroxymethyl) aminomethane (Tris) (cat # 252859) and ethylenediaminetetraacetic acid (EDTA) (cat # E5134) were purchased from Sigma-Aldrich. Bovine serum albumin (BSA) (cat # MB083) was purchased from HiMedia. Hemin was dissolved in DMSO in 1 mM concentration as a stock solution and serially diluted to 20 µM concentration as a working solution in citrate phosphate buffer pH 4.5. Hydrogen peroxide solution (cat # Q18755), 30% was purchased from Qualigens. Streptavidin-magnetic bead was purchased from Merck. Personal glucometer such as Dr. Morepen GLuco One (model no: BG-03, licence no: MFG/IVD/2021/000034 and lot no: AIK237) and Apollo pharmacy blood glucose monitoring (model no: APG01, licence no: MFG/IVD/2021/000034 and lot no: DIL272) and their manufacturer designated strips were purchased locally and was used as it is for all analysis experiments. All experiments utilized type I water. All concentrations below are final concentrations in the actual process or reaction unless stated otherwise.

**Oligonucleotide sequences:** Biotinylated oligonucleotide sequence, 5'-[biotin] AAA AAA AAA ACT TTG ACT TCA TAG GAT CCA TGG TAA GCC-3' underlined region is "sequence a*"; Inactive DNAzyme sequence: 5'-*CTG GGA GGG AGG GAG GG*A AAA AAA GGC TTA CCA TGG ATC CTA TGA A-3' (underlined region is "sequence a", that would bind the biotinylated sequence's "sequence a*", italics region would convert to EAD2 DNAzyme[1]). Biotinylated oligonucleotide sequence and inactive DNAzyme were purchased from Merck (HPLC purification) and Eurofin (HPSF purification), respectively.

**Construction of DNAzyme-hemin complex:** Binder-DNAzyme sequence (5 µM) was added to a G-quadruplex assembly buffer (Tris-HCl (12.5 mM, pH 7.6), 75 mM of NaCl, 10 mM KCl, 0.015% Triton-X-100, and 0.5% DMSO), heated at 95°C for 5 min, slowly cooled in a dry bath, and then stored at 4°C. For hemin incorporation, the inactive DNAzyme sequence (1 µM) in the same buffer was mixed with hemin (2 µM) and incubation was done for 1 h at room temperature in the dark.

**Immobilization of DNAzyme into beads:** 16.67 µg of streptavidin-coated magnetic beads were washed 3 times with $TEN_{100}$ buffer (10 mM Tris-HCl, 100 mM NaCl, 1 mM EDTA, pH-7.5). 25 picomoles of biotinylated oligonucleotide sequence (final concentration 0.5 µM) in 50 µL $TEN_{100}$ was added to the streptavidin-coated magnetic beads and incubated for 20 minutes at room temperature. Following 2 rounds of magnetic decantation wash with $TEN_{100}$ buffer, blocking was carried out by incubating magnetic beads with 1% BSA (bovine serum albumin) in 50 µL $TEN_{100}$ buffer for 20 minutes. Following 2 rounds of magnetic decantation wash with $TEN_{1000}$ buffer (10 mM Tris-HCl, 1M NaCl, 1 mM EDTA pH-7.5) and 2 round of G-quadruplex assembly buffer wash, 25 picomoles of DNAzyme-hemin complex (final concentration 500 nM) in 50 µL in 1X G-quadruplex assembly buffer was incubated with biotinylated oligonucleotide-conjugated streptavidin magnetic beads for 20 minutes. After 2 rounds of magnetic decantation wash with G-quadruplex assembly buffer, the DNAzyme-hemin complex (500 nM) immobilized into streptavidin magnetic beads via biotinylated oligonucleotide was stored in 50 µL of G-quadruplex assembly buffer at 4°C. To analyze the activity of immobilized DNAzyme-hemin complex, a colorimetric test was carried out each time before subjecting it to spectrophotometric or glucometer assays. ABTS (2,2'-azino-bis(3-ethylbenzothiazoline-6-sulfonic acid)) 1.8 mM), hydrogen peroxide ($H_2O_2$) 50 mM), and citrate-phosphate buffer (pH 5.0) were added to the DNAyme-hemin complex (50 nM and 12.5 nM) bound to the magnetic beads. The color would swiftly appear within 2-3 min, proving continued activity of DNAzyme.

**Generation of ferricyanide concentration vs absorbance vs glucometer standard curve:** An experiment was conducted for comparing two different methods for measuring the concentration of potassium ferricyanide: glucometer readings and absorbance measurements (ELISA reader). Different ferricyanide concentration (1, 2.5, 5, 10, 20, 40, 60, 80, 100, 120, and 140 mM) where prepared in MES buffer pH 5.0. Absorbance (405 nm) and glucometer readings (mg/dL) were recorded for each concentration.

**Optimization of various buffers to evaluate DNAzyme catalytic activity:** The buffers employed for the reactions included MES (pH 5.0), sodium acetate (pH 5.0), Tris-HCl (pH 7.6), Tris-HCl (pH 8.5), HEPES (pH 7.2), HEPES (pH 8.2). Each buffer was used at a final concentration of 12.5 mM supplemented with salts NaCl (final 75 mM) and KCl (final 10 mM). Two experimental conditions were evaluated for each buffer type: DNAzyme-positive (DNAzyme +) and DNAzyme-negative (DNAzyme −). For each condition and buffer, 100 µl of reaction mixture was prepared in replicates, containing the buffer, Na and K salts (concentrations as above), potassium ferrocyanide [$K_4Fe(CN)_6$] (10 mM), $H_2O_2$ (50 mM), and, for the DNAzyme-positive condition only, DNAzyme (final 50 nM, unless mentioned otherwise) was used. 100 µl of each reaction mixture was added to a 96-well plate in three or more replicates. Absorbance at 405 nm was measured using a microplate reader. The kinetic loop program was set in the plate reader to take absorbance readings at 5-minute intervals, starting from 0 minutes and continuing through to 40 minutes (i.e., at 0, 5, 10, 15, 20, 25, 30, 35, and 40 minutes), without removing the plate from the reader during the entire assay. The entire reaction was carried out at room temperature. Additionally, glucometer readings were recorded using a standard digital glucometer (Dr. Morepen GLuco One) under both experimental conditions at 5-minute intervals over a total duration of 40 minutes. For each time point, triplicate readings were taken for all buffers by adding

1-2 µL of reaction mixture onto the test strip. The glucometer reading, reported in mg/dL, reflected the extent of oxidation of potassium ferrocyanide to ferricyanide in presence and absence of DNAzyme.

**Optimization of $H_2O_2$ concentration for DNAzyme catalytic activity:** The reactions were conducted in either MES (12.5 mM, pH 5.0) or HEPES (12.5 mM, pH 7.2) buffer. For each condition and different $H_2O_2$ concentration, 100 µl of reaction mixture were prepared in three or more replicates. Each mixture contained the respective buffer (MES pH 5.0 or HEPES pH 7.2, 12.5 mM), salts (NaCl and KCl as above), 10 mM potassium ferrocyanide, the specified concentration of $H_2O_2$, and, for DNAzyme-positive condition only, 50 nM DNAzyme. The 100 µl reaction was added to wells of a 96-well plate, with each $H_2O_2$ concentration tested under both DNAzyme-absent and DNAzyme-present conditions. Absorbance at 405 nm was measured at room temperature using a plate reader. Readings were recorded every 5 minutes over a total duration of 40 minutes using the kinetic loop function. This setup allowed the real-time monitoring of the oxidation of potassium ferrocyanide catalyzed by the DNAzyme in the presence of various $H_2O_2$ concentrations. Additionally, glucometer readings were taken for reactions containing 5 mM and 10 mM $H_2O_2$ for both MES pH 5.0 and HEPES pH 7.2 as mentioned above.

**Optimization of potassium ferrocyanide concentration:** These reactions were performed using varying concentrations of ferrocyanide (1 mM, 2.5 mM, 4 mM, 6 mM, 8 mM, 16 mM, 25 mM, and 50 mM) in the presence of two optimized $H_2O_2$ concentrations (5 mM and 10 mM). For each condition and ferrocyanide concentration, 100 µl of reaction mixture was prepared in replicates. Each mixture contained MES buffer (12.5 mM, pH 5.0), salts (NaCl and KCl as above), the specified concentration of potassium ferrocyanide, $H_2O_2$, (either 5 mM or 10 mM), and (for DNAzyme-positive condition) DNAzyme (50 nM) was used. Absorbance was measured at 405 nm starting from 0 minutes and continuing at 5-minute intervals up to 45 minutes (i.e., at 5, 10, 15, 20, 25, 30, 35, and 40 minutes), without removing the plate from the reader during the entire assay.

**Determination of optimal DNAzyme-hemin concentration range:** These reactions were conducted in MES buffer (12.5 mM, pH 5.0), salts (NaCl and KCl as above), $H_2O_2$ (5 mM or 10 mM, as specified), ferrocyanide (concentrations as specified in "Results" section), and DNAzyme (as specified in Results). For each reaction condition, 100 µl of reaction mixture was prepared in replicates. All reactions were carried out at room temperature. The catalytic activity was monitored by measuring absorbance at 405 nM using a microplate reader set to kinetic loop mode. Absorbance readings were recorded at multiple time points: 0, 5, 10, 15, 20, 25, 30, 35, and 40 minutes. Also, glucometer readings were done for the same at all different time points for both DNAzyme-absent and DNAzyne-present condition.

**Glucometer measurement**: The mixture of the reagents was same as mentioned above. Majority of the experiments were carried out using "Dr. Morepen GLuco One brand", while investigation with an alternate brand was performed with “Apollo pharmacy blood glucose monitoring” brand. At pre-specified time point, 1 μL of the reaction was aliquoted onto a hydrophobic parafilm strip. The glucometer measurement was then carried out as instructed in the manufacturer’s manual using the 1 μL droplet. In the case of Dr. Morepen GLuco One or Apollo pharmacy blood glucose monitoring, it involved first inserting the strip electrode into the glucometer, waiting for a blood droplet image to appear on screen (within 2-3 sec), and then touching the reaction droplet with respective electrode at the designated place for blood droplet insertion.

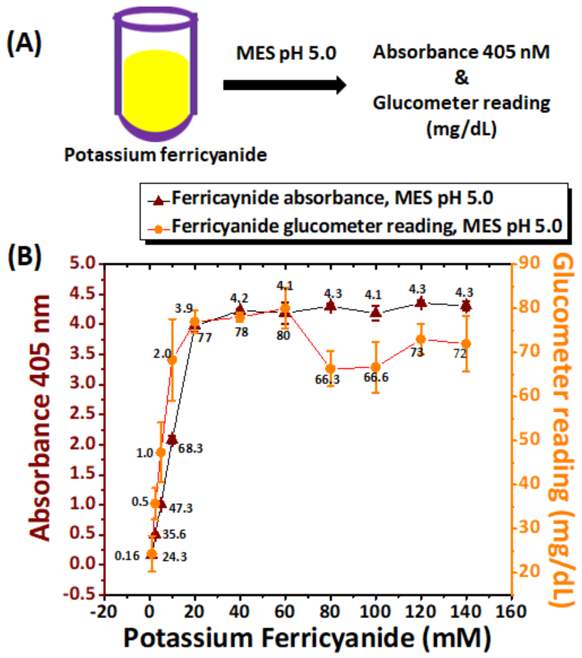


**Figure S1.** Correlation between ferricyanide concentration and glucometer readings in MES buffer. The graph shows the relationship between ferricyanide concentration (x-axis) and the two measurement methods: absorbance at 405 nm (y-axis, left) and glucometer readings in mg/dL (y-axis, right). Glucometer measurements were carried out in Dr. Morepen GLuco One brand

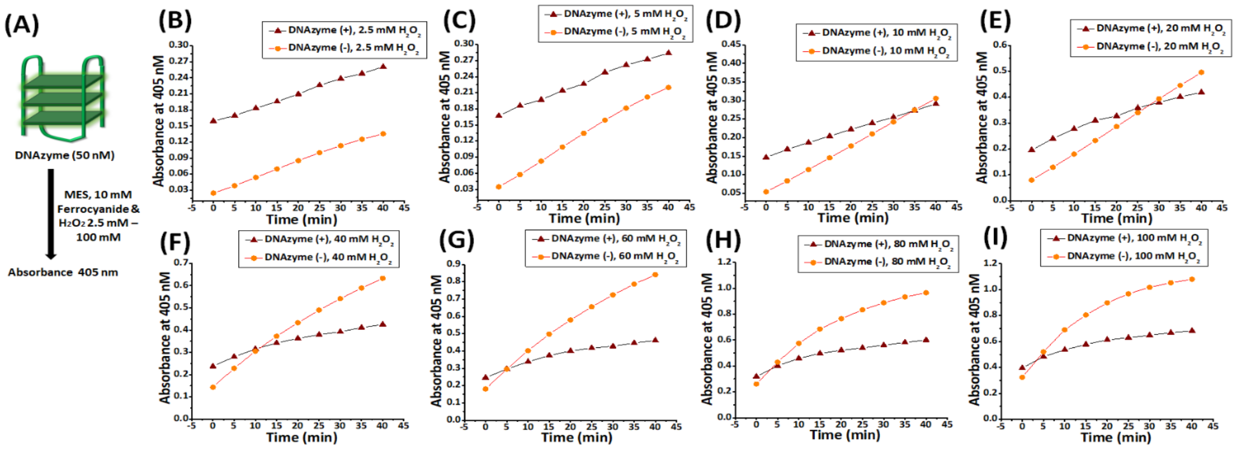


**Figure S2.** Optimization of $H_2O_2$ concentration for DNAzyme-catalyzed oxidation of potassium ferrocyanide (10 mM) to ferricyanide. Panel A illustrates the schematic representation of the experimental setup. Panel B-I show the time dependent change in absorbance from 0 to 40 minutes measured in the presence or absence of 50 nM DNAzyme, with MES buffer at (pH 5.0), 10 mM potassium ferrocyanide, and varying $H_2O_2$ concentrations: 2.5 mM (Panel B), 5 mM (Panel C), 10 mM (Panel D), 20 mM (Panel E), 40 mM (Panel F), 60 mM (Panel G), 80 mM (Panel H), and 100 mM (Panel I).

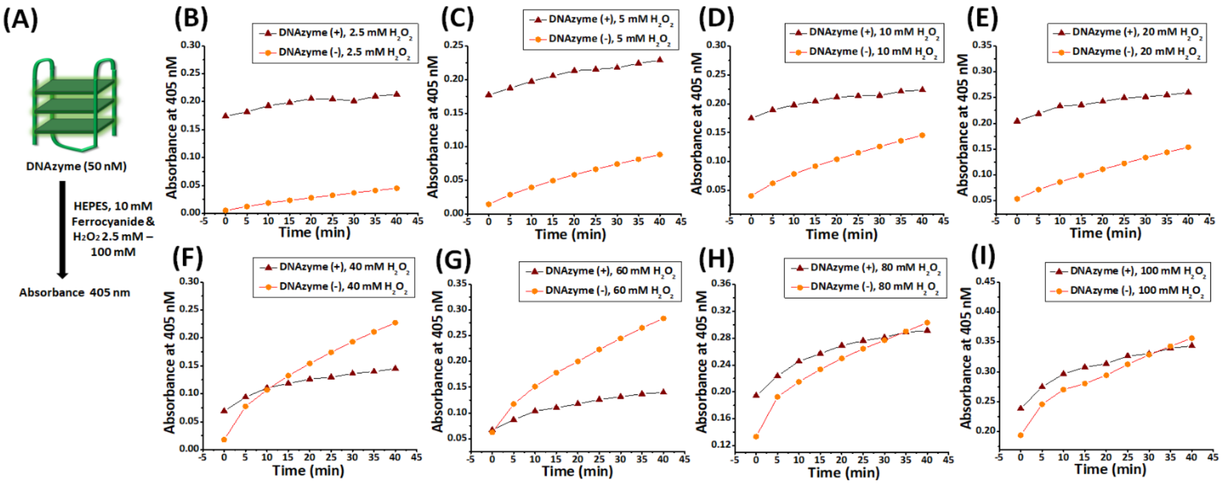


**Figure S3.** Optimization of $H_2O_2$ concentration for DNAzyme-catalyzed oxidation of potassium ferrocyanide (10 mM) to ferricyanide. Panel A illustrates the schematic representation of the experimental setup. Panel B-I show the representative time dependent change in absorbance from 0 to 40 minutes measured in the presence or absence of 50 nM DNAzyme, with HEPES buffer at (pH 7.2), 10 mM potassium ferrocyanide, and varying $H_2O_2$ concentrations: 2.5 mM (Panel B), 5 mM (Panel C), 10 mM (Panel D), 20 mM (Panel E), 40 mM (Panel F), 60 mM (Panel G), 80 mM (Panel H), and 100 mM (Panel I).

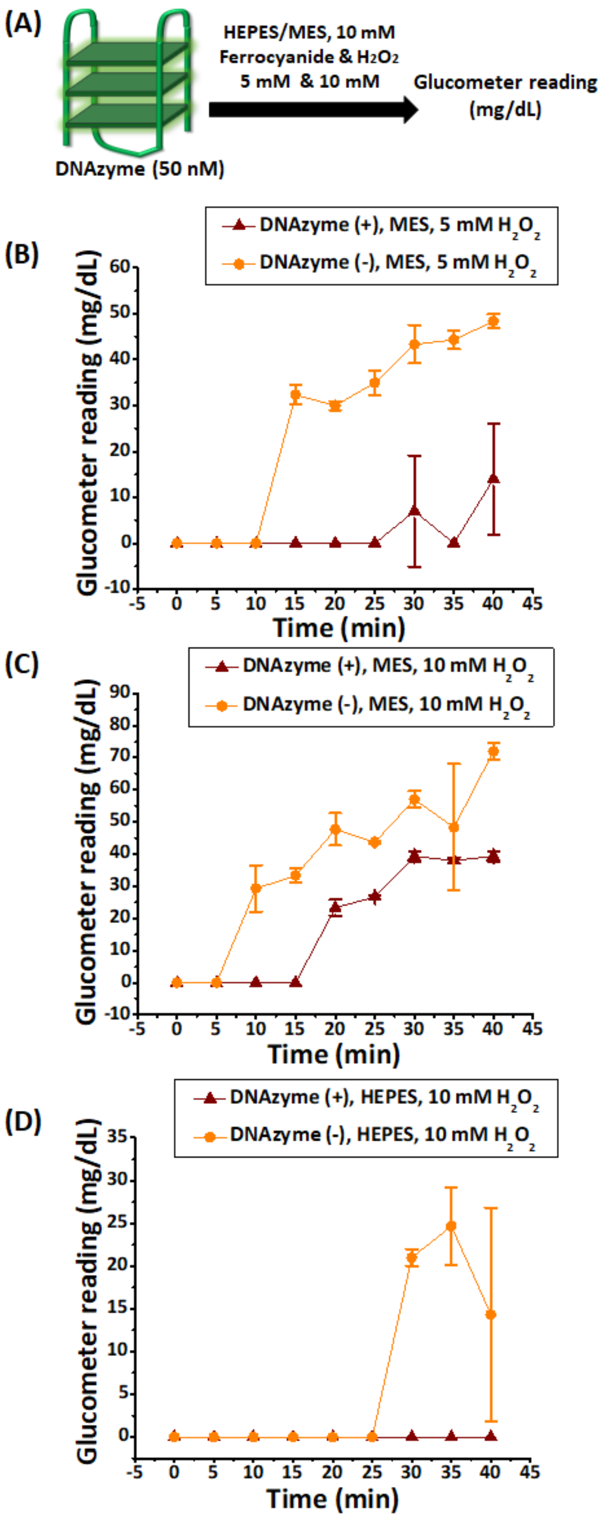


**Figure S4.** Effect of varying $H_2O_2$ concentration buffer on DNAzyme-catalyzed oxidation of potassium ferrocyanide (10 mM) to ferricyanide monitored by glucometer. Panel B-D represents the representative time course of glucometer readings (mg/dL) from 0 minutes to 40 minutes, measured at 5 minutes intervals, in different experimental conditions: 5 mM $H_2O_2$ in MES buffer (pH 5.0) (Panel B) and 10 mM of $H_2O_2$ in MES (pH 5.0) (Panel C) and HEPES (pH 7.2) (Panel D) buffers, each containing 10 mM ferrocyanide in the presence and absence of DNAzyme. Experiments were performed with Dr. Morepen GLuco One brand. The zero indicates absence of a measurable value.

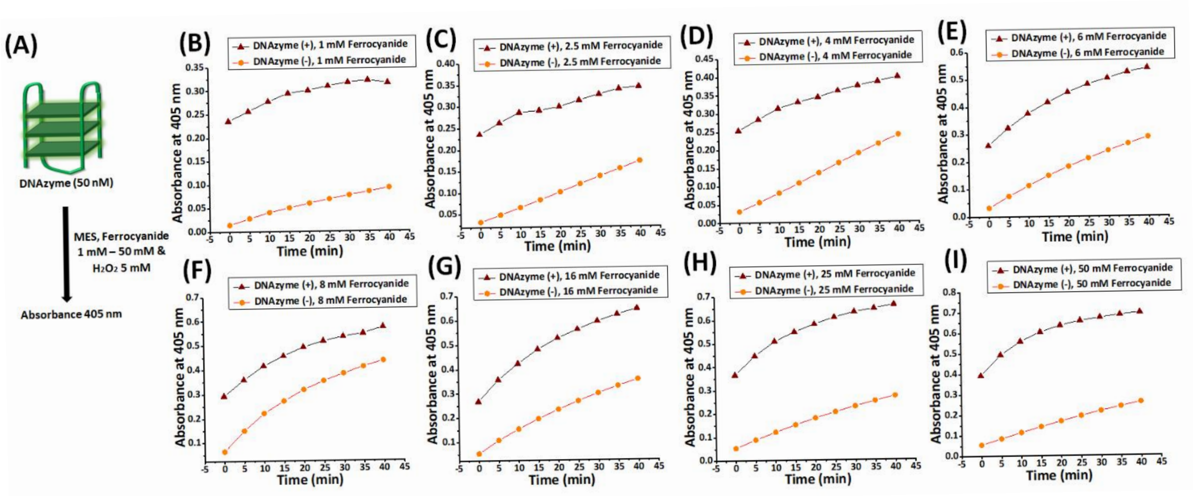


**Figure S5.** Optimization of ferrocyanide concentration on DNAzyme-catalysed oxidation of potassium ferrocyanide to ferricyanide in 5 mM $H_2O_2$. Panel A illustrates the schematic representation of the experimental setup. Panel B-I show the representative time dependent change in absorbance from 0 to 40 minutes measured in the presence or absence of 50 nM DNAzyme, with MES buffer at (pH 5.0), 5 mM of $H_2O_2$, and varying ferrocyanide concentrations: 1 mM (Panel B), 2.5 mM (Panel C), 4 mM (Panel D), 6 mM (Panel E), 8 mM (Panel F), 16 mM (Panel G), 25 mM (Panel H), and 50 mM (Panel I).

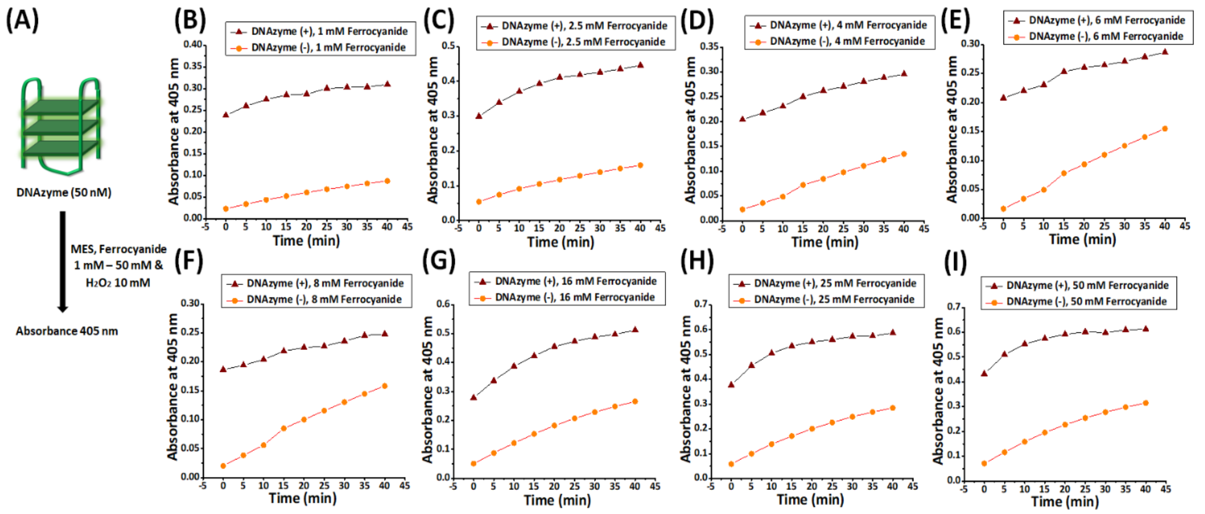


**Figure S6.** Optimization of ferrocyanide concentration on DNAzyme-catalysed oxidation of potassium ferrocyanide to ferricyanide in 10 mM $H_2O_2$. Panel A illustrates the schematic representation of the experimental setup. Panel B-I show the representative time dependent change in absorbance from 0 to 40 minutes measured in the presence or absence of 50 nM DNAzyme, with MES buffer at (pH 5.0), 10 mM of $H_2O_2$, and varying ferrocyanide concentrations: 1 mM (Panel B), 2.5 mM (Panel C), 4 mM (Panel D), 6 mM (Panel E), 8 mM (Panel F), 16 mM (Panel G), 25 mM (Panel H), and 50 mM (Panel I).

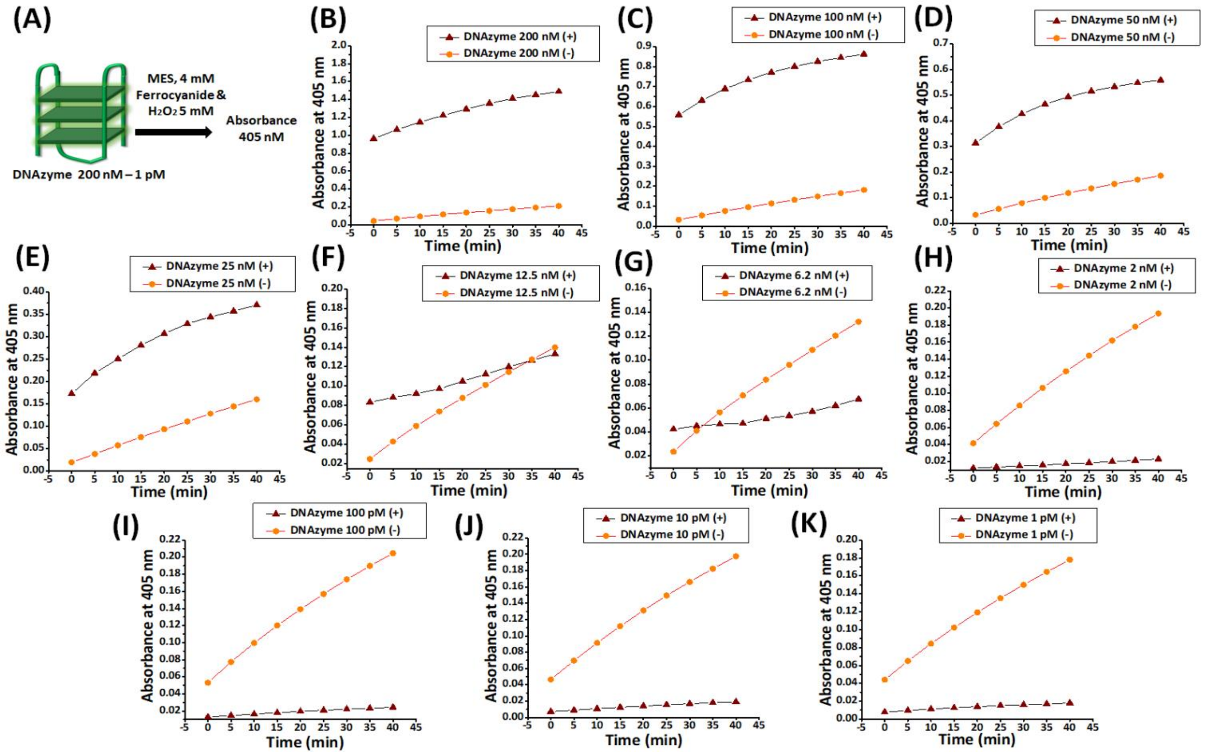


**Figure S7.** Optimization of DNAzyme concentration range for DNAzyme-catalyzed oxidation of potassium ferrocyanide to ferricyanide (4 mM) in 5 mM $H_2O_2$. Panel A provides a schematic representation of the experimental setup. Panel B-K shows the representative time-dependent change in absorbance from 0 to 40 minutes measured in MES buffer (pH 5.0), with a constant 5 mM of $H_2O_2$, 4 mM ferrocyanide concentration, in the absence and presence of DNAzyme with varying DNAzyme concentrations: 200 nM (Panel B), 100 nM (Panel C), 50 nM (Panel D), 25 nM (Panel E), 12.5 nM (Panel F), 6.2 nM (Panel G), 2 nM (Panel H), 100 pM (Panel I), 10 pM (Panel J), and 1 pM (Panel K).

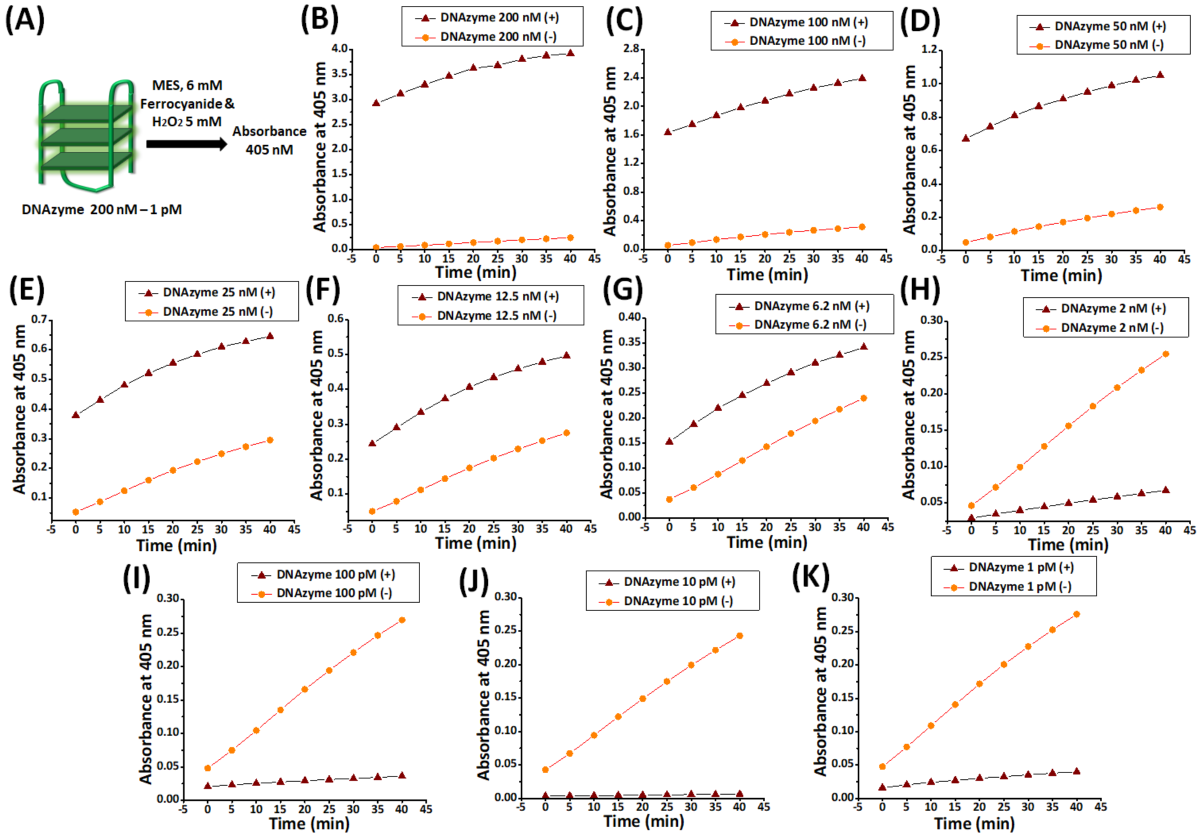


**Figure S8.** Optimization of DNAzyme concentration range for DNAzyme-catalyzed oxidation of potassium ferrocyanide to ferricyanide (6 mM) in 5 mM $H_2O_2$. Panel A provides a schematic representation of the experimental setup. Panel B-K shows the representative time-dependent change in absorbance from 0 to 40 minutes measured in MES buffer (pH 5.0), with a constant 5 mM of $H_2O_2$, 6 mM ferrocyanide concentration, in the absence and presence of DNAzyme with varying DNAzyme concentrations: 200 nM (Panel B), 100 nM (Panel C), 50 nM (Panel D), 25 nM (Panel E), 12.5 nM (Panel F), 6.2 nM (Panel G), 2 nM (Panel H), 100 pM (Panel I), 10 pM (Panel J), and 1 pM (Panel K).

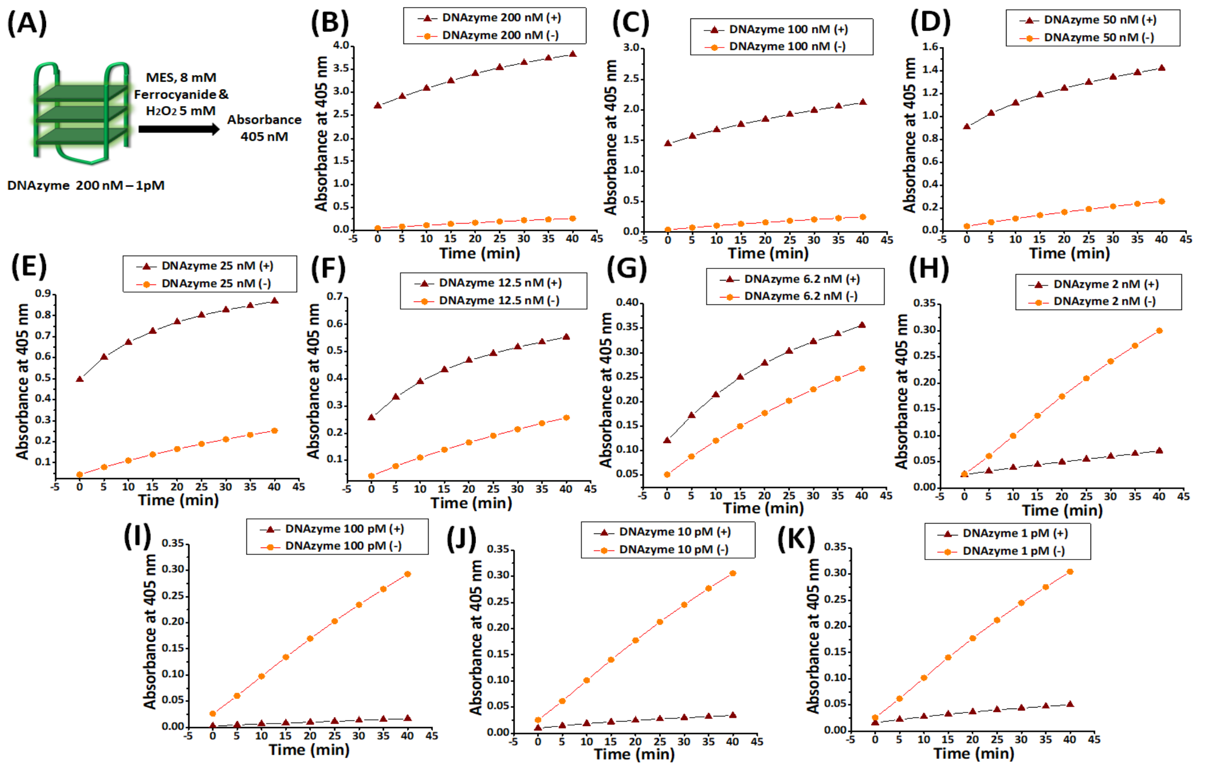


**Figure S9.** Optimization of DNAzyme concentration range for DNAzyme-catalyzed oxidation of potassium ferrocyanide to ferricyanide (8 mM) in 5 mM $H_2O_2$. Panel A provides a schematic representation of the experimental setup. Panel B-K shows the representative time-dependent change in absorbance from 0 to 40 minutes measured in MES buffer (pH 5.0), with a constant 5 mM of $H_2O_2$, 8 mM ferrocyanide concentration, in the absence and presence of DNAzyme with varying DNAzyme concentrations: 200 nM (Panel B), 100 nM (Panel C), 50 nM (Panel D), 25 nM (Panel E), 12.5 nM (Panel F), 6.2 nM (Panel G), 2 nM (Panel H), 100 pM (Panel I), 10 pM (Panel J), and 1 pM (Panel K).

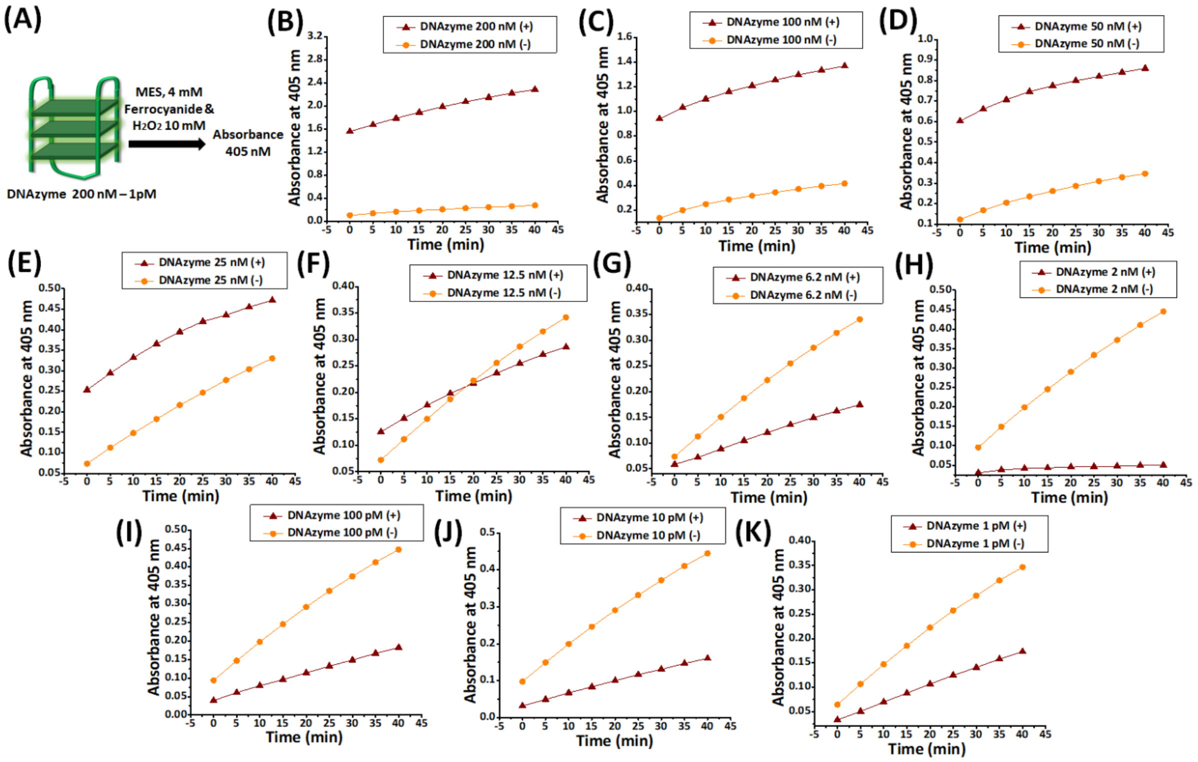


**Figure S10.** Optimization of DNAzyme concentration range for DNAzyme-catalyzed oxidation of potassium ferrocyanide to ferricyanide (4 mM) in 10 mM $H_2O_2$. Panel A provides a schematic representation of the experimental setup. Panel B-K shows the representative time-dependent change in absorbance from 0 to 40 minutes measured in MES buffer (pH 5.0), with a constant 10 mM of $H_2O_2$, 4 mM ferrocyanide concentration, in the absence and presence of DNAzyme with varying DNAzyme concentrations: 200 nM (Panel B), 100 nM (Panel C), 50 nM (Panel D), 25 nM (Panel E), 12.5 nM (Panel F), 6.2 nM (Panel G), 2 nM (Panel H), 100 pM (Panel I), 10 pM (Panel J), and 1 pM (Panel K).

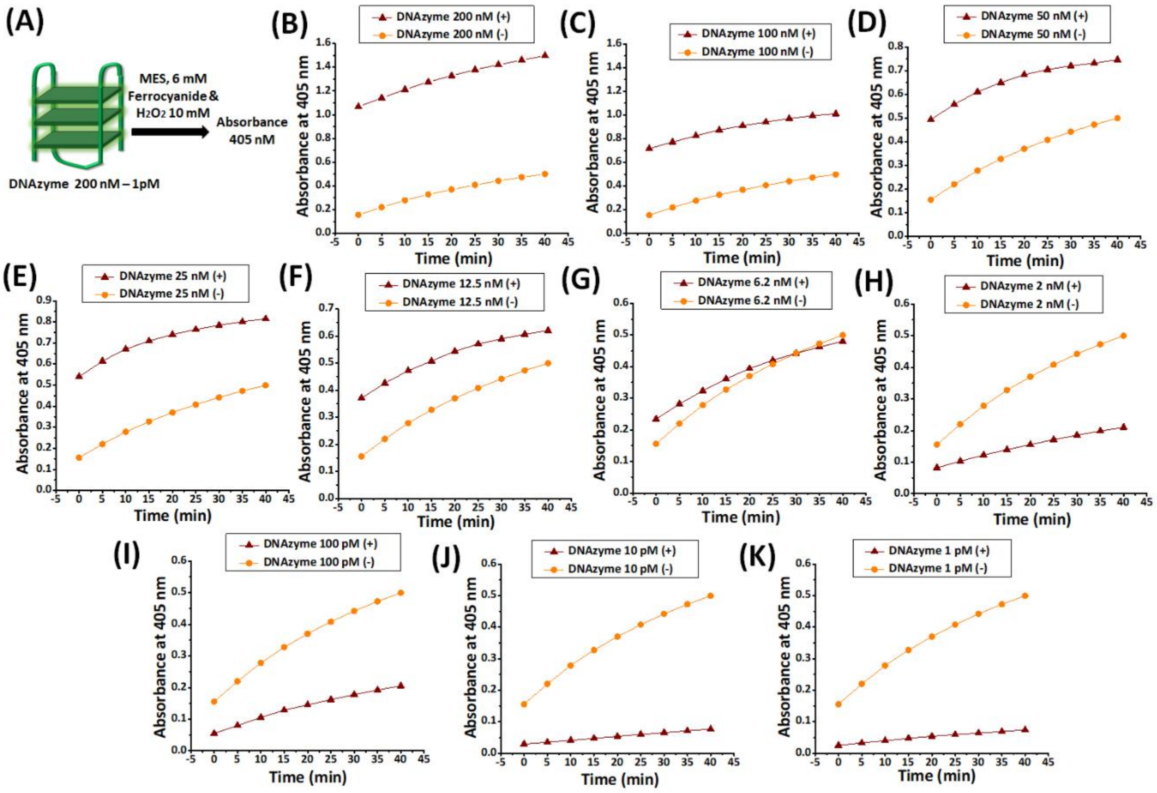


**Figure S11.** Optimization of DNAzyme concentration range for DNAzyme-catalyzed oxidation of potassium ferrocyanide (6 mM) to ferricyanide in 10 mM $H_2O_2$. Panel A provides a schematic representation of the experimental setup. Panel B-K shows the representative time-dependent change in absorbance from 0 to 40 minutes measured in MES buffer (pH 5.0), with a constant 10 mM of $H_2O_2$, 6 mM ferrocyanide concentration, in the absence and presence of DNAzyme with varying DNAzyme concentrations: 200 nM (Panel B), 100 nM (Panel C), 50 nM (Panel D), 25 nM (Panel E), 12.5 nM (Panel F), 6.2 nM (Panel G), 2 nM (Panel H), 100 pM (Panel I), 10 pM (Panel J), and 1 pM (Panel K).

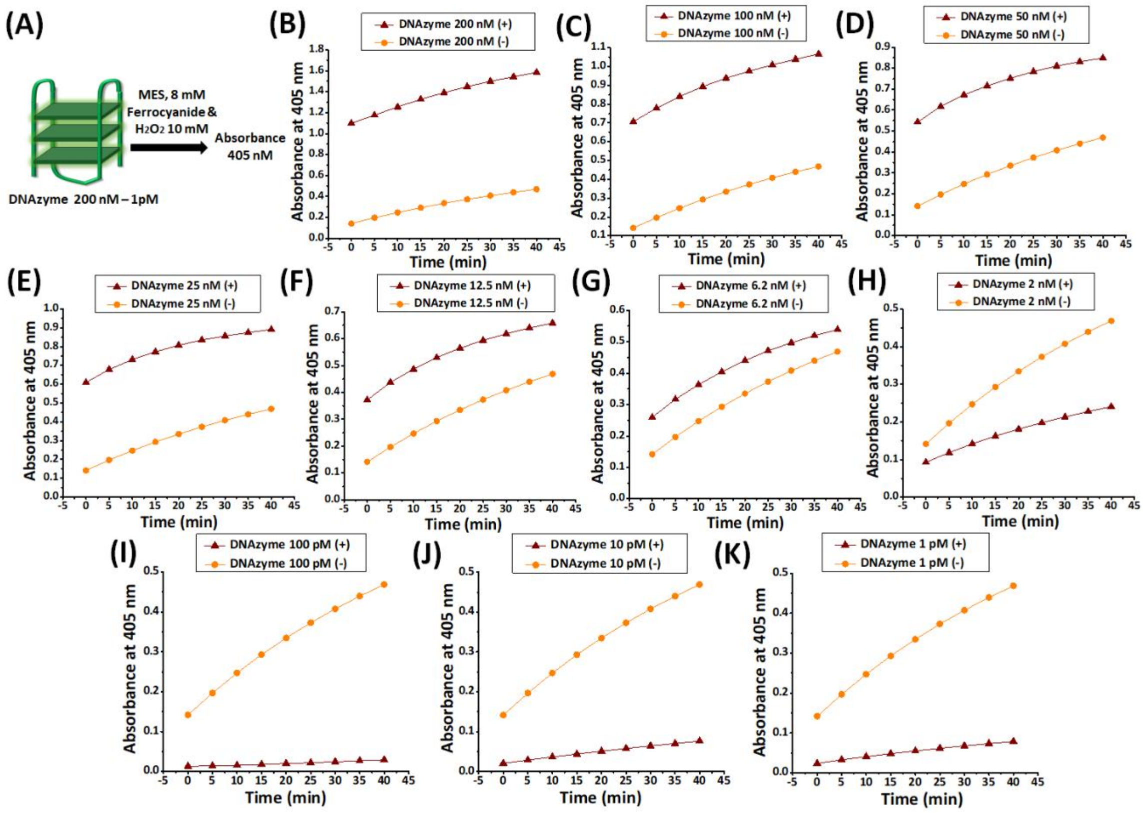


**Figure S12.** Optimization of DNAzyme concentration range for DNAzyme-catalyzed oxidation of potassium ferrocyanide (8 mM) to ferricyanide in 10 mM $H_2O_2$. Panel A provides a schematic representation of the experimental setup. Panel B-K shows the representative time-dependent change in absorbance from 0 to 40 minutes measured in MES buffer (pH 5.0), with a constant 10 mM of $H_2O_2$, 8 mM ferrocyanide concentration, in the absence and presence of DNAzyme with varying DNAzyme concentrations: 200 nM (Panel B), 100 nM (Panel C), 50 nM (Panel D), 25 nM (Panel E), 12.5 nM (Panel F), 6.2 nM (Panel G), 2 nM (Panel H), 100 pM (Panel I), 10 pM (Panel J), and 1 pM (Panel K).

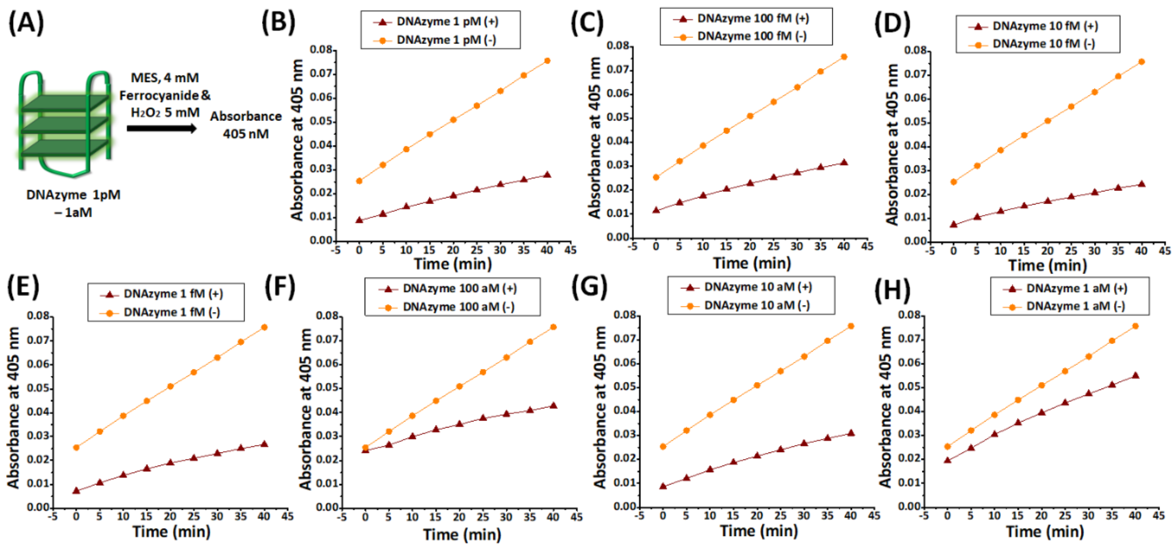


**Figure S13.** Optimization of lower DNAzyme concentration range for DNAzyme-catalyzed oxidation of potassium ferrocyanide (4 mM) to ferricyanide in $H_2O_2$. Panel A provides a schematic representation of the experimental setup. Panel B-H shows the representative time-dependent change in absorbance from 0 to 40 minutes measured in MES buffer (pH 5.0), with a constant 5 mM of $H_2O_2$, 4 mM ferrocyanide concentration, in the absence and presence of DNAzyme with lower DNAzyme concentrations: 1 pM (Panel B), 100 fM (Panel C), 10 fM (Panel D), 1 fM (Panel E), 100 aM (Panel F), 10 aM (Panel G), and 1 aM (Panel H).
